# Diversity of function and higher-order structure within HWE sensor histidine kinases

**DOI:** 10.1101/2020.07.08.194225

**Authors:** Igor Dikiy, Danielle Swingle, Kaitlyn Toy, Uthama R. Edupuganti, Giomar Rivera-Cancel, Kevin H. Gardner

## Abstract

Integral to the protein structure/function paradigm, oligomeric state is typically conserved along with function across evolution. However, notable exceptions such as the hemoglobins show how evolution can alter oligomerization to enable new regulatory mechanisms. Here we examine this linkage in histidine kinases (HKs), a large class of widely distributed prokaryotic environmental sensors. While the majority of HKs are transmembrane homodimers, members of the HWE/HisKA2 family can deviate from this architecture as exemplified by our finding of a monomeric soluble HWE/HisKA2 HK (EL346, a photosensing Light-Oxygen-Voltage (LOV)-HK). To further explore the diversity of oligomerization states and regulation within this family, we biophysically and biochemically characterized multiple EL346 homologs and found a range of HK oligomeric states and functions. Three LOV-HK homologs are primarily dimeric with differing structural and functional responses to light, while two Per-ARNT-Sim (PAS)-HKs interconvert between differentially active monomers and dimers, suggesting dimerization might control enzymatic activity for these proteins. Finally, we examined putative interfaces in a dimeric LOV-HK, finding that multiple regions contribute to dimerization. Our findings suggest the potential for novel regulatory modes and oligomeric states beyond those traditionally defined for this important family of environmental sensors.

## Introduction

Sensor histidine kinases (HKs) are signal transduction receptors that are widespread in prokaryotes, enabling these organisms to sense and respond to diverse environmental stimuli (3, 4). HKs are typically found in two-component systems (TCS), which are most simply composed of a sensor HK and a downstream response regulator (RR) co-located in the same operon. The RR responds to a signal from the HK by acting as a phosphorylation-dependent transcription regulator, with activation often triggering RR DNA binding and altering the expression of nearby genes (5-8).

The prototypical sensor HK contains N-terminal sensor and C-terminal kinase domains, the latter of which can be subdivided into a two-helix dimerization histidine phosphotransfer (DHp) domain and a catalytic ATP-binding (CA) domain (9). The detection of a stimulus by the sensor domain (or in some cases, elsewhere (8)) modulates the CA domain autophosphorylation of a conserved His residue in the DHp domain; the phosphate group in this phospho-His adduct is subsequently transferred to a conserved Asp in a downstream RR (5).

Most characterized sensor HKs are obligate homodimeric transmembrane receptors, facilitating the transmission of signals across the membrane from periplasmic sensor domains to cytoplasmic kinase domains via symmetry-breaking conformational changes (9-12). The regulation of typical sensor HKs is largely believed to require a dimeric complex, both for the structural changes between the “off” and “on” state as well as determining whether autophosphorylation occurs between protomers (*in trans*) or within a protomer (*in cis*) (9, 13). Dimerization is mediated via the DHp domains of two monomers, forming a four-helix bundle and facilitating pivot-, piston-, or scissoring-type movements as part of the activation process (9, 14), echoing similar themes in integrins (15) and certain other transmembrane receptors. In contrast, some sensors adopt different oligomeric states, such as the large multimeric assemblies of chemotaxis HKs (16), hexameric KaiC circadian HKs (17) or the monomeric photosensory EL346 HK (1, 18).

The traditional view of sensor HK structure has been built up from extensive structural and biochemical studies (9) that have usually focused on HKs in only one of several subfamilies of histidine kinases (the HisKA family within Pfam (19)). In contrast, the related HWE (His-Trp-Glu)-HK and HisKA_2 families (referred to here as the HWE/HisKA2 family) of HKs have been less studied from a structural perspective (14). Members of this group, which contain specific sequence variations in the DHp and CA domains compared to members of the canonical HisKA family, tend to signal to a more diverse set of output proteins which often lack DNA-binding output domains (14). The HWE/HisKA2 family is also enriched in HKs lacking transmembrane segments, likely localizing them to the cytosol. Notably, soluble sensor HKs are relieved of the constraints of transmitting a signal across a lipid bilayer, which typically involves signal transmission via motions of one protomer with respect to the other (9, 14). This raises several questions: Are such soluble HKs able to adopt diverse oligomeric states, including non-dimeric architectures? And could these varied architectures be equivalently varied in their regulation? We view the relatively understudied HWE/HisKA2 family of HKs (14) as a suitable testbed for these questions.

Our prior work on the HWE/HisKA2 family focused on three soluble HKs involved in the general stress response of the marine alphaproteobacterium *Erythrobacter litoralis* HTCC2594 (EL346, EL362, and EL368) (1, 18, 20, 21). Each of these proteins senses blue light through a light oxygen voltage (LOV) sensor domain (22, 23), a sub-class of the versatile Per-ARNT-Sim (PAS) domains that are present in over 30% of HKs (4, 24). The response to blue light provides a readily tractable way to explore the inactive and active states of histidine kinases, typically corresponding to dark and light states of LOV-HKs (21, 25, 26). These states can be conveniently distinguished spectrophotometrically: while the dark state spectra of LOV-HKs exhibit a triplet peak at around 450 nm, characteristic of protein-bound flavin chromophores, the light state spectrum has a broad, singlet absorption peak at 390 nm (27). This spectroscopic change is due to the photochemically-triggered reduction of the flavin chromophore and concomitant formation of a cysteine-C(4a) covalent adduct, a bond that is thermally reversible in the dark on a timescale varying from seconds to hours among LOV domains (23).

While our prior studies indicate that EL368 is dimeric, EL346 is a soluble and functional monomer (1, 20). The EL346 crystal structure reveals that it is held in a monomeric state by an intramolecular association between its DHp-like domain and the LOV sensory domain, blocking the face typically used for DHp domains to dimerize (1). Despite this deviation from current HK signaling models, the monomeric EL346 undergoes light-induced increases in autophosphorylation rate (1) as well as conformational changes involving the sensor, DHp, and kinase domains (18). More broadly, the active monomer of EL346 suggests that soluble HKs may not require dimerization as a key element in signal transduction, unlike conventional transmembrane HKs.

To address this general issue in HK signaling, we set out to characterize the oligomeric and functional states of several novel HWE/HisKA2 HKs. To do so, we used sequence similarity searches to identify multiple soluble LOV- and PAS-HKs closely related to EL346, selecting five proteins for in-depth analyses. We showed that three of these five are novel dimeric LOV-HKs, each of which properly photocycles by UV-visible absorbance spectroscopy. Notably, these LOV-HKs differed in the effect of light on their autophosphorylation activity, with a mix of dark- and light-activated proteins among them. In addition, we also characterized two light-insensitive PAS-HKs which equilibrate between monomeric and dimeric states with differing activities, suggesting dimerization as a possible regulatory switch in this family of HKs. Finally, we combined structural and sequence information to engineer a “monomerized dimer” by deleting putative dimerization interfaces in a dimeric LOV-HK, finding that broadly distributed residues contribute to dimerization. Taken together, our data reveal a wider range of oligomerization states and functional activity in HKs than traditionally described.

## Results

### Bioinformatics analysis reveals numerous alphaproteobacterial EL346 homologs

To identify other potential monomeric histidine kinases, we searched for enzymes highly similar in domain organization to the well-validated and functional EL346 monomer. In addition to EL346, the top 100 BLASTp hits defined a set of proteins from *Alphaproteobacteria* which all contained the same domain architecture of a single N-terminal PAS or LOV domain followed by a C-terminal HWE/HisKA2 histidine kinase. None of these sequences contained any predicted transmembrane segments, strongly suggesting all these candidates are soluble. A multiple sequence alignment of these protein sequences and distance tree analysis revealed three clusters (**Fig. 1a**), which split on increasingly diverse sensor types: EL346-like LOV-HKs, LOV-HKs, and PAS-HKs (non-LOV). LOV domains were provisionally identified by the presence of the characteristic “NCRFLQ” sequence and high homology (> 30% identity) to the EL346 LOV domain (28), while the non-LOV PAS domains lacked this sequence motif (highlighted in **Fig. S1**). Interestingly, a subgroup of the EL346-like LOV-HKs is missing the conserved phosphoacceptor histidine (**Fig. S1**). We also included the previously-identified related EL368 (dimer) and EL362 (mixed oligomerization state) histidine kinases from *E. litoralis* HTCC2594 (20, 21), as well as the *Caulobacter crescentus* LovK (dimer) (29) in the alignment; all three of these proteins grouped in the LOV-HKs cluster. Additionally, our search identified a third PAS-HK cluster with proteins containing non-LOV PAS domains that shared 17-24% sequence identity with the EL346 LOV domain. Notably, many of these domains contained a conserved tryptophan in place of the cysteine utilized in LOV-type photochemistry (28) (**Fig. S1**). These data show that the LOV/PAS-HK architecture is widespread throughout *Alphaproteobacteria* (as are HWE/HisKA2 HKs in general (30, 31)) and contains members which could well respond to stimuli other than blue light.

**Figure 1:**
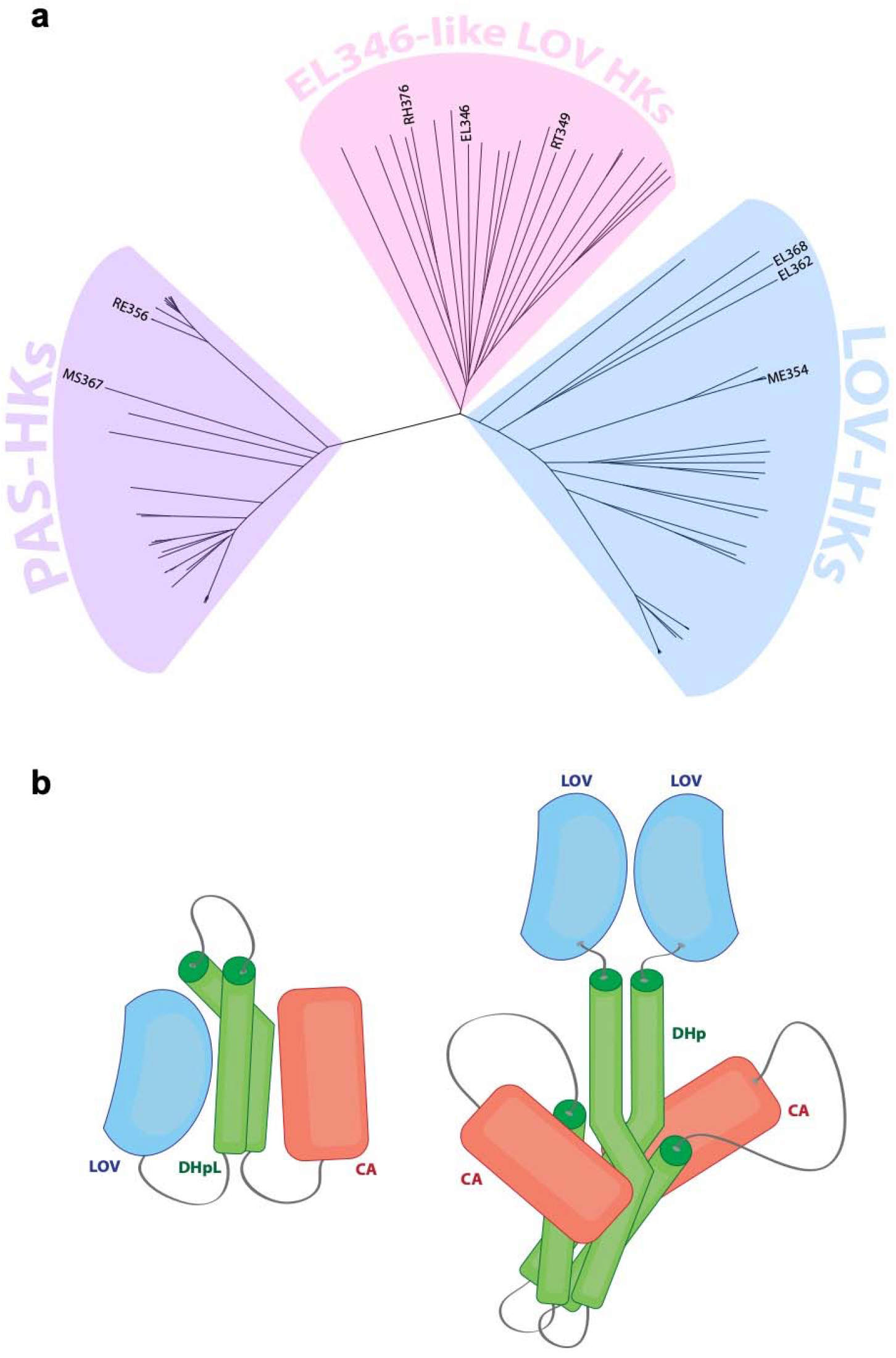
EL346 homologs form three distinct clusters. **a)** PAS-HKs (purple), EL346-like LOV HKs (pink), and LOV-HKs (blue). The names of the proteins from each cluster that were selected for this study are shown. All other names are omitted for simplicity and can be found in **Figure S2**. **b)** Schematic structures of monomeric (left) and dimeric (right) LOV-HKs in the dark state. Monomeric schematic is based on the structure of EL346 (1) and dimeric is based on the engineered fusion protein YF1 (2).

### LOV homologs are light sensing and appear mostly dimeric, but have diverse functional responses to illumination

We posited that the EL346-like LOV-HK cluster might contain a mix of other monomeric and dimeric HKs, respectively analogous to EL346 and most natural HKs (as well as the engineered LOV-HK YF1 (2)), as illustrated in **Fig. 1b**. We selected several HKs from each cluster and characterized their oligomeric state and function with a variety of biochemical approaches. We started by determining the ability of the LOV-containing proteins to bind flavin chromophores and undergo typical LOV photochemistry via UV-visible absorbance spectroscopy. The absorbance spectra of dark and light state samples of four representative proteins – EL346, ME354, RT349, and RH376 (see **Table 1** for protein details) – all showed the characteristic flavin-LOV triplet (∼450 nm) in the dark state which disappeared upon illumination (**Fig. 2a**). Upon incubation in the dark, the 450 nm absorbance returned with first-order exponential kinetics that varied by protein: the LOV-HKs measured here, such as ME354 (236 min), RT349 (240 min), and RH376 (62 min), had longer reversion constants *τ* than EL346 (33 min) (**Fig. 2b**). The particularly long reversion time constants of ME354 and RT349 can be rationalized by the presence of several “slowing” mutations in their LOV domains relative to that of EL346, specifically at positions 19, 21, 32, and 101 (EL346 numbering), as collated in a review by Pudasaini and coworkers (**Fig S3**) (23).

**Figure 2:**
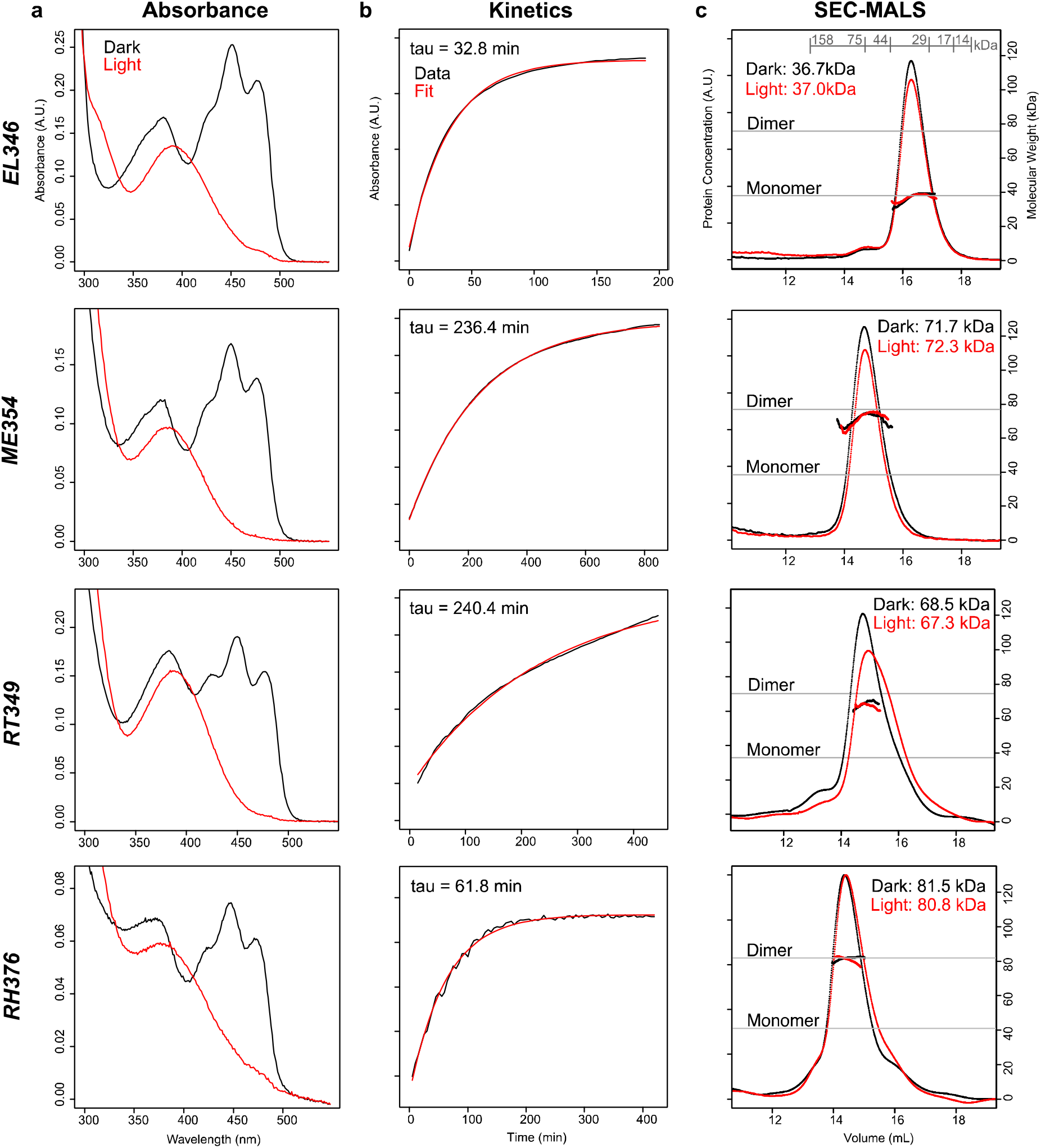
UV-visible absorption spectra and SEC-MALS reveal diversity of LOV-HK dark state reversion kinetics and oligomerization states. **a)** UV-visible absorption spectra of EL346, ME354, RT349, and RH376, superimposing dark state scans (black) with scans post-illumination (red). All spectra display the disappearance of the 450 nm triplet upon exposure to light, consistent with LOV photochemistry. Spectra were recorded in the presence of 1 mM ATP. **b)** Dark state reversion kinetics, measured by the return of absorbance at 446 nm post illumination (black), fit to single exponentials (red) reveal an 8-fold range of time constants. **c)** Superdex 200 GL SEC-MALS chromatograms are represented by dRI in arbitrary units and MALS-derived masses are solid lines under the peaks (n=1). MWs and elution volumes of six standard proteins are shown in grey in the top panel. MALS traces show that EL346 is monomeric in solution, while ME354, RT349 and RH376 are all predominantly dimeric (Table 1). A light-dependent shift in elution peak shape and position is seen in RT349.

**Table 1:**
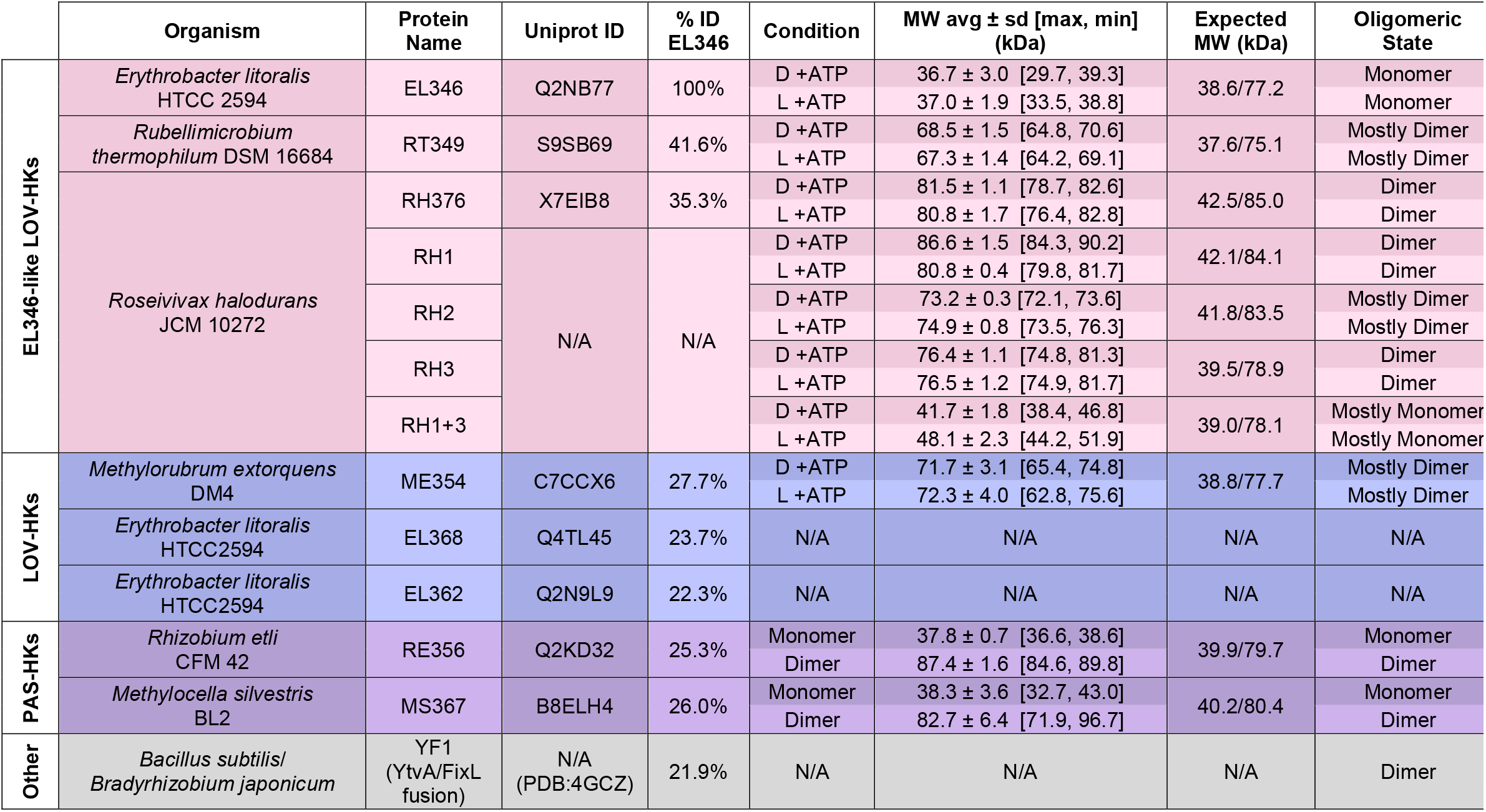
Oligomeric states of known LOV-HKs and PAS-HKs include mix of monomers and dimers. Soluble HKs shown here are separated by cluster with the same coloring as Fig. 1 (EL346-like LOV HKs – pink, LOV-HKs – blue, PAS-HKs – purple, Other – grey) and include those investigated in this study (EL346, RT349, RH376, ME354, RE356, and MS367) as well as those identified in literature (EL368, EL362, and YF1) (2, 20). Mass data were collected using Superdex200 GL SEC-MALS (n=1). Values shown are MALS MW (average mass), minimum, maximum, and standard deviation among the points that make up the mass line for each measurement, and oligomeric states were defined using a qualitative five-bin scale.

We determined the oligomeric state of these LOV-HKs in the presence of ATP (1 mM) in dark and light conditions using size exclusion chromatography coupled to multi-angle light scattering (SEC-MALS). We began by confirming that EL346 is monomeric under both conditions with minimal change in global shape upon illumination as revealed by SEC elution profiles. In contrast, the three other LOV-HK proteins were dimeric or mostly dimeric, with RT349 exhibiting a light-dependent shift in elution peak position and shape (**Fig. 2c, Table 1**). Further investigation showed that the RT349 elution profile was also impacted by ATP as well: in the absence of ATP, we observed two new peaks at even higher elution volumes in dark and light conditions (**Fig. S4**).

While the oligomeric states of the three new LOV-HKs were all similar, analyses of autophosphorylation activity indicated a diverse activity profile (**Fig. 3**). EL346 and RH376 exhibited the light-dependent increase in activity typically seen in LOV-HKs, albeit to different extents. In contrast, RT349 displayed an inverted signaling logic, with higher autophosphorylation rates in the dark state than in the light state. While this phenomenon has been reported for engineered LOV-HKs such as YF1 variants (2) and the EL346 R175A mutant (1), it does not seem to have been observed in a naturally-occurring LOV-HK. On the other end of the spectrum, ME354 exhibited no detectable activity in both the dark and light, suggesting that even though this protein properly photocycles, its kinase activity is not regulated directly by light under these conditions. Overall, our results illustrate that despite adopting similar oligomeric states, the LOV-HKs studied here exhibit diverse autophosphorylation activities, suggesting varying forms of regulation and different signaling paradigms.

**Figure 3:**
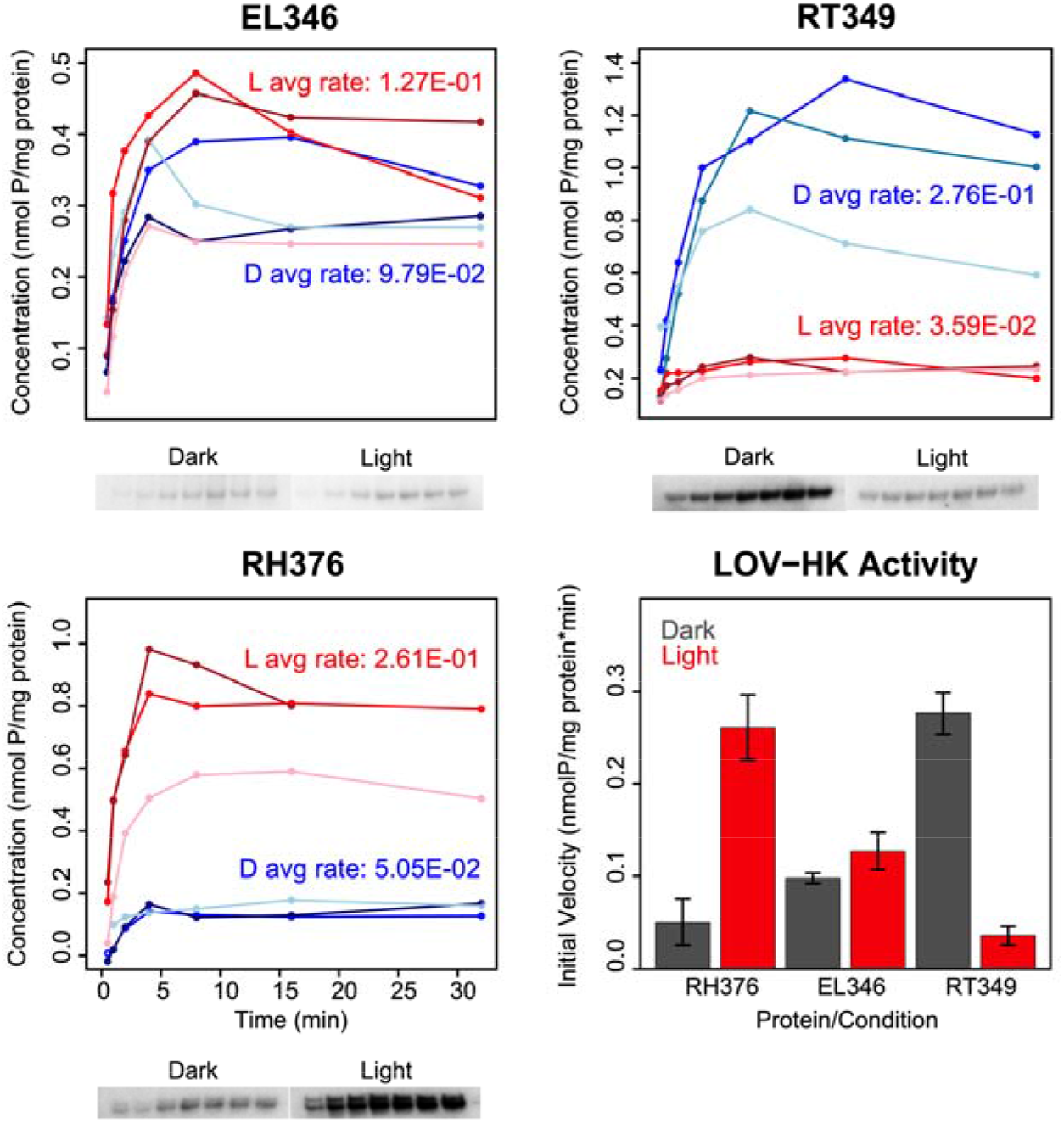
LOV-HKs display differing activity levels. Autophosphorylation assays (plots, above; phosphor images, below) of EL346 and RH376 follow the typical trend for LOV-HKs found in nature, with light activation increasing activity levels, while RT349 produces opposite results: it is more active in the dark state than in the light. Each bar indicates the mean +/- one standard deviation (n=3).

### Non-LOV PAS-HKs contain mixture of monomers and dimers, with each state exhibiting different activity levels and stabilities

The two PAS-HKs characterized here, from *Rhizobium etli* CFN 42 (RE356) and from *Methylocella silvestris* BL2 (MS367), exhibit 45% sequence identity overall to one another (32% over the PAS domain). In SEC-MALS experiments, both proteins appeared as mixtures of dimer and monomer, as well as an oligomeric or aggregated fraction. We found that the monomeric and dimeric states of MS367 and RE356 were in slow equilibrium, allowing them to be separately purified and re-injected onto SEC-MALS (**Fig. 4a**). We characterized the *in vitro* stability of the two different states by incubating mixed samples at 4*°*C for varying amounts of time and using size-exclusion chromatography to visualize elution peaks. For both PAS-HKs, we observed that the monomeric state was stable on the order of days, while the dimer slowly oligomerized or aggregated on this timescale (**Fig. 4b**).

**Figure 4:**
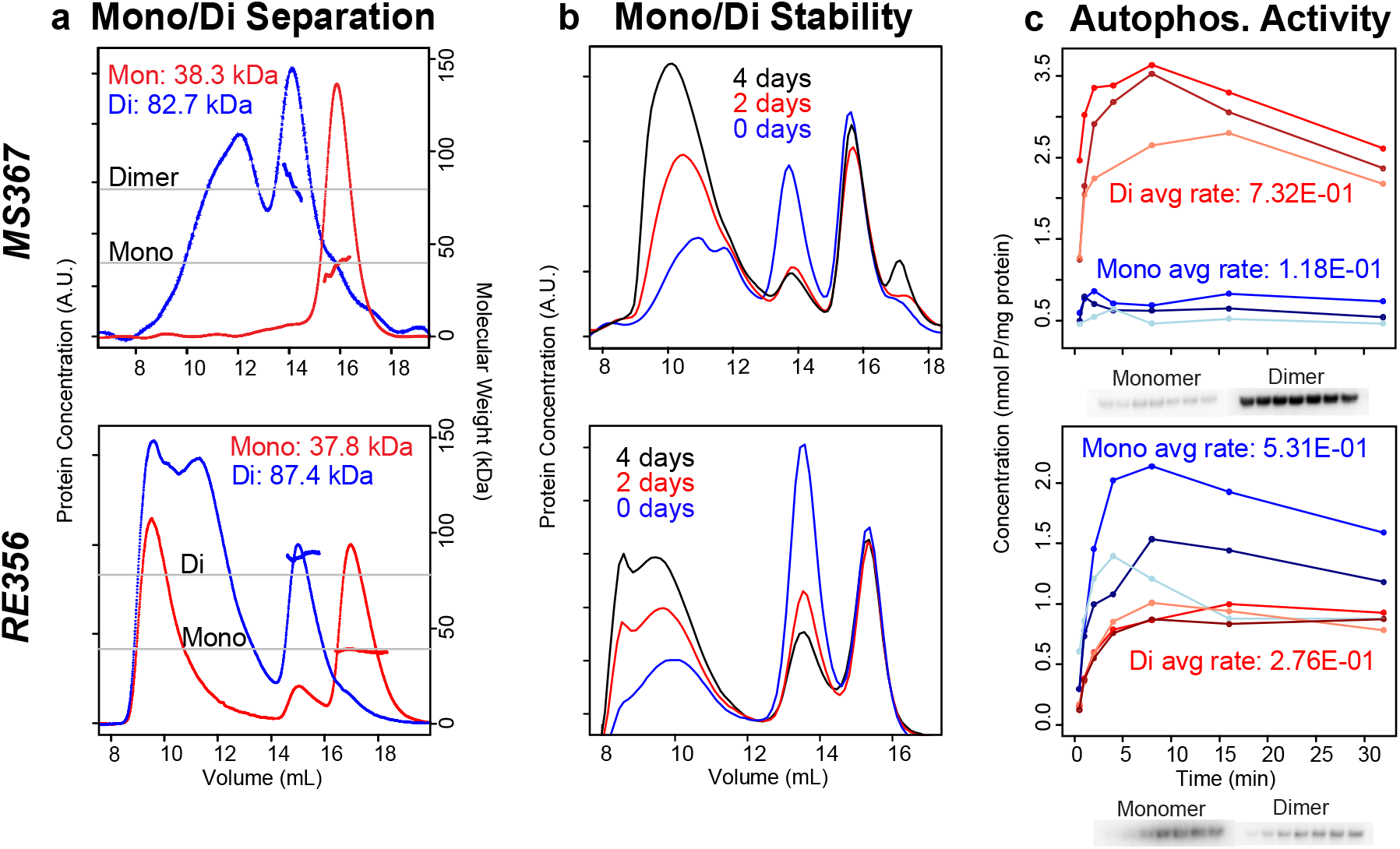
The PAS-HKs MS367 and RE356 are monomer/dimer mixtures that display differential autophosphorylation activity and stabilities. **a)** SEC-MALS measurements of PAS-HKs MS367 and RE356 reveal the presence of both monomeric and dimeric states. Monomers (red traces) and dimers (blue traces) were separated and injected onto a Superdex200 GL SEC (n=1). The resulting MALS-derived masses (solid lines under elution peaks) show that, under these conditions, both states remain stable upon separation. **b)** Superdex200 GL SEC traces of PAS-HK monomer/dimer mixtures after being stored at 4 °C for 0, 2, and 4 days at ∼20 µM concentration, with both proteins exhibiting a similar trend of the monomer remaining stable over time, while the dimer tends to aggregate. **c)** Autophosphorylation assays (n=3) (plots, above; phosphor images, below) of the monomer (blue traces) and dimer (red traces) of the PAS-HKs show that for MS367, the dimer has a higher rate of activity while for RE356, the monomer has a higher rate.

Though these proteins currently lack a known trigger analogous to light for the LOV-HKs, we took advantage of the slow equilibrium between oligomeric states to determine whether they exhibited differing levels of autophosphorylation. We observed that both MS367 and RE356 displayed autophosphorylation activity in their monomeric and dimeric states with the characteristic plateau indicating an equilibrium between kinase and phosphatase activity (**Fig. 4c**). While the MS367 dimer appears to be more active than the monomer, RE356 produced opposite results, with the monomer displaying more activity than the dimer. Despite similarity in primary sequence and oligomeric assembly, the two PAS-HKs studied here in an *in vitro* context have apparently opposite regulation via dimerization, again illustrating the diversity of function across this family.

### Engineering a “monomerized” LOV-HK

Next, we sought to identify the determinants of dimerization in this family of sensor HKs, recognizing that sequence changes can influence the multimeric arrangement of proteins belonging to the same family (32, 33). To guide our analysis, we aligned the sequence of the monomeric EL346 with those of selected dimeric HKs from all three clusters. These pairwise alignments suggested four candidate regions that might influence dimerization; three regions missing from EL346, but found in many of the other HKs (named RH1-3), as well a unique C-terminal segment in EL346 (RH4) (**Figs. 5a and S5**).

**Figure 5:**
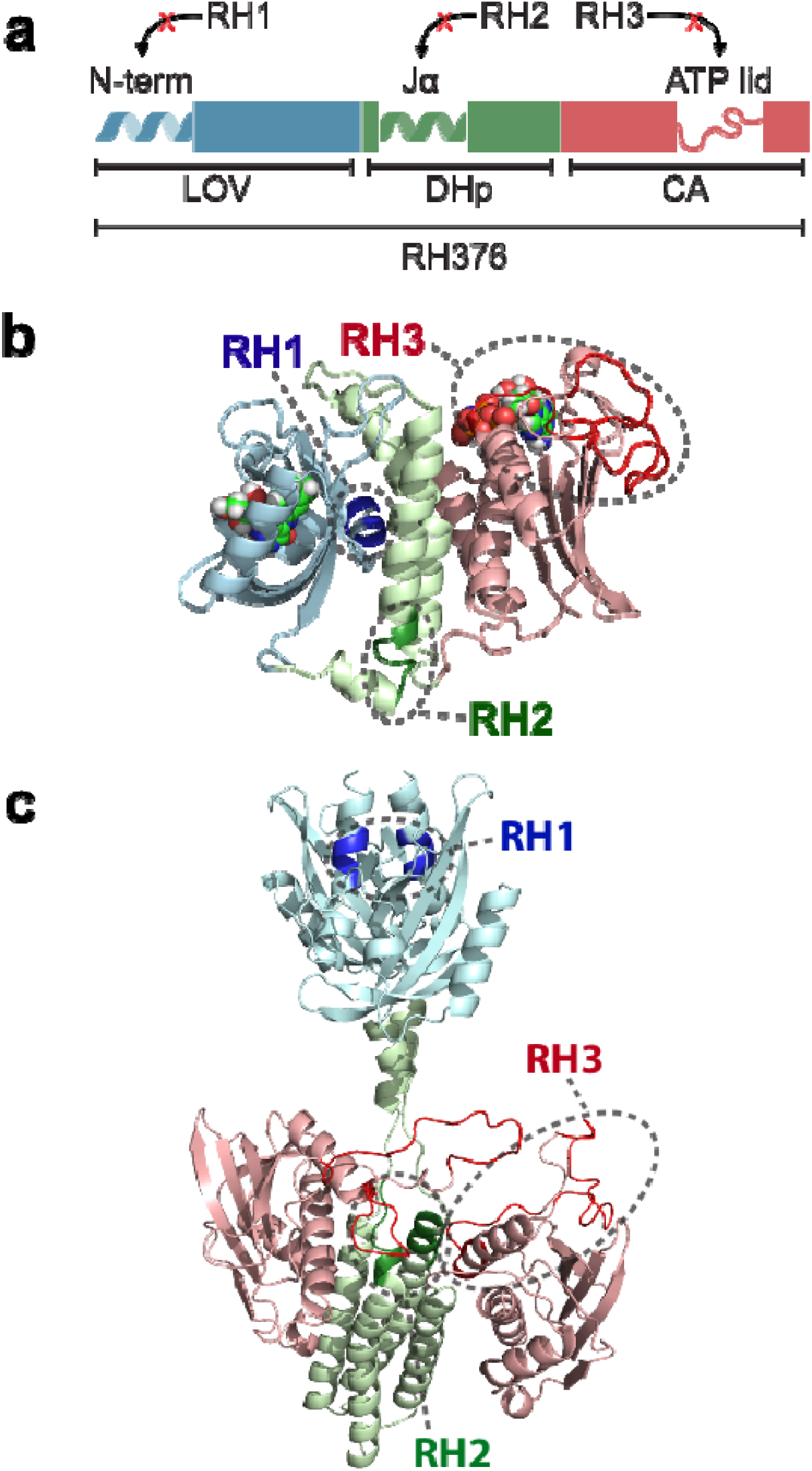
RH376 mutant deletion regions are designed in predicted dimer interfaces. RH376 deletion mutants RH1, RH2, and RH3 are highlighted in 3 schematics: **a)** Domain architecture with the predicted secondary structure at the location of each deletion. RH1 is located in the A’α helix of the LOV domain, RH2 in the linker between the LOV domain and α1 of the DHp domain, and RH3 is in the ATP lid. **b)** Monomeric homology model of RH376 based on the structure of EL346 (1). **c)** Dimeric homology model of RH376 based on the engineered YF1 sensor kinase (2). All LOV domains are colored blue, DHp domains green, and CA domains red. Mutations are highlighted and circled.

To test the hypothesis that one or more of the RH1-4 regions might be determinants of oligomeric state, we systematically deleted one or more of these regions from the dimeric RH376 protein, which contained all four segments, in the hope of finding a constitutively monomeric variant (**Figs. 5b and 5c**). As the RH4 segment corresponds almost entirely to the highly dynamic C-terminal residues we previously deleted without effect on the monomeric state of EL346 (18), we did not investigate RH4 as a driver of monomer/dimer state and instead focused on the RH1-3 segments unique to RH376.

Observing that the insert regions were either in previously-reported dimerization interfaces for LOV domains (RH1, A’*α* helix) (2, 20, 34) and HKs (RH2, DHp domain) (35, 36), or large insertions with predicted secondary structure elements (RH3, ATP lid), we hypothesized that these segments could contribute to the dimerization of RH376 and perhaps other HKs. A RH376 homology model (37) based on the engineered dimeric LOV-HK YF1 protein (2) illustrates the potential involvement of these residues in dimerization interfaces (**Fig. 5c**). We thus deleted each of the RH1, RH2, and RH3 regions from the RH376 sequence both individually and in combination, expressed these variant proteins, and compared them to RH376 WT in terms of LOV photocycle, oligomeric state, and autophosphorylation activity.

RH1, RH2, and RH3 proteins displayed the characteristic LOV absorbance triplet centered at 450 nm and exhibited dark state reversion kinetics similar to the WT protein (**Fig. S6**), suggesting that the deletions did not perturb the folding and function of the LOV domain. In RH2, we detected a small change in the relative heights of the triplet peaks, indicating that flavin binding is likely adversely affected (**Fig. S6**) despite the deletion occurring outside the LOV domain. We determined the oligomeric state of each deletion mutant by SEC-MALS; while none of the single deletion constructs eluted as a monomer regardless of dark or light conditions, RH2 had an altered measured mass and SEC elution volume which suggest it samples both dimeric and monomeric conformations (**Table 1, Fig. 6**).

**Figure 6:**
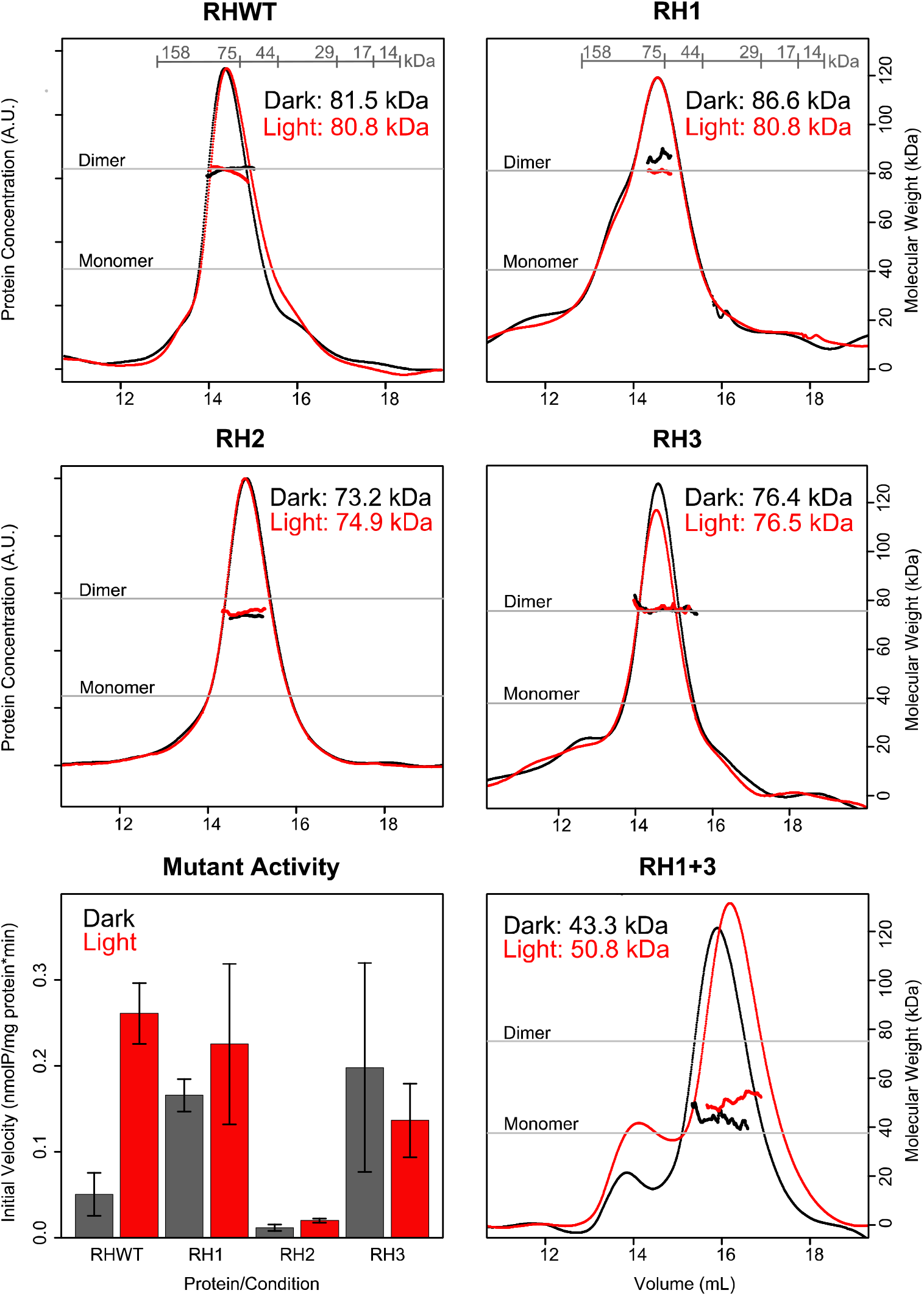
RH376 mutants RH1, RH2 and RH3 are active and mostly dimeric, while double mutant RH1+3 is inactive and primarily monomeric. Superdex 200 GL SEC-MALS results (n=1) show that RH376 WT, RH1 and RH3 are clear dimers, while RH1+3 is primarily monomeric, and RH2 is primarily dimeric. Expected MWs (grey lines) were calculated based on protein sequence. Mutant activity, represented by initial velocity calculated from the linear part of the curve in at least three replicates for each protein/condition (bars represent average), is shown in the bottom left panel. No activity was detected for RH1+3. Light activation can be seen in RH376 WT, RH1 and RH2, but not RH3. Each bar indicates the mean +/- one standard deviation (n=3).

We suspected that none of these deletions would have a major impact on autophosphorylation activity of RH376, given their modest effects on photocycle and oligomerization state. As a baseline, we found that the RH376 WT protein displays robust autophosphorylation activity in the dark as well as a ∼5x increase in initial velocity upon illumination (**Table S1, Fig. 6**). The RH1 mutant showed autophosphorylation activity similar to the WT in the light, but greater in the dark, resulting in a much lower degree of light-activation. Similarly, RH2 retained some light activation, but showed very little activity in both dark and light conditions (**Table S1, Fig. 6**). On the other hand, RH3 displayed activity in the dark that was ∼4x the WT initial velocity, with a light-dependent decrease in activity (**Table S1, Fig. 6**). In all, each mutant had a markedly different effect on autophosphorylation activity despite having minimal effects on dimerization.

Given that neither RH1 nor RH3 perturbed dimeric state and both displayed activity levels comparable to WT, yet had divergent effects on light-activation (**Table S1, Fig. 6**), we explored the effects of combining these deletions (“RH1+3”). While neither the LOV absorbance spectrum nor photocycle of this protein were affected (**Fig. S6**), SEC-MALS showed a dramatic change in oligomeric state, as RH1+3 existed mainly as a monomer, though neither deletion alone showed monomeric character (**Table S1, Fig. 6**). Additionally, we detected no autophosphorylation activity in RH1+3 at conditions similar to the other proteins tested. Overall, we conclude that RH376 uses a distributed set of residues to control oligomeric state.

## Discussion

Our work further expands the known behavior of sensor HKs to include two proteins in slow monomer:dimer equilibria along with three chiefly dimeric proteins, one of which exhibits a slight conformational change upon activation, and all of which exhibit varying activities. Prior to this study, the vast majority of characterized sensor HKs were dimers with the EL346 LOV-HK from *E. litoralis* HTCC2594 (1, 20) as the only well-characterized, functional protein that we are aware of as an exclusive monomer. Using sequence homology searches to find homologs of EL346, we found a large family of related LOV- and PAS-HKs in the *Alphaproteobacteria*. Most of these proteins were Pfam (19) HisKA_2 or HWE-HK-type histidine kinases, two groups of HKs that have been less well-characterized structurally to date (1, 12, 14, 38), and we hypothesized that a subset of these sequences with the highest homology to monomeric EL346 might contain other monomers.

Contrary to this expectation, we found a diverse range of assemblies, conformational changes, and autophosphorylation activity among the five new sensor HKs characterized here. For the LOV-HK proteins, we found several dimers, including two members of the EL346-like cluster of LOV-HKs (RT349 and RH376) which SEC-MALS analyses showed to be dimeric or primarily dimeric in both dark and light conditions. ME354, a newly characterized member of the other LOV-HK cluster is also mostly dimeric, similar to the established EL368 and *C. crescentus* LovK (20, 29), which both also cluster within that second group.

While these new LOV-HKs are all mostly dimeric, we observed differences in their structural response to blue light activation as assessed by SEC-MALS: ME354 and RH376 showed minimal light-dependent change, while RT349 eluted at later volumes in the light, suggesting that it adopts a more compact structure under this condition. Notably, this shift was dependent upon not only light, but also the presence of ATP (**Fig. S4**). These light-dependent changes are more substantial than we previously observed by SEC-MALS with the EL346 and EL368 LOV-HKs (1, 18, 20), and are perhaps similar to the kinds of changes seen by small angle solution X-ray scattering in in the engineered YF1 protein (2, 39). Further studies are needed to more fully characterize these motions and establish how they relate to the large conformational changes postulated to accompany activation from crystal structures of inactive HKs (40, 41). RT349 is also very distinctive in that it is more active in the dark than in the light **(Fig. 3).** While this behavior has been observed before in variants of engineered (2) and mutants of natural LOV-HKs (1), we are not aware of any naturally-occurring LOV-HKs that share this signaling polarity. We anticipate that such a light-dependent change in net autophosphorylation has contributions from altered kinase and phosphatase activities (42), and this will be the focus of a subsequent investigation.

Turning to the two PAS-HKs, both RE356 and MS367 interconvert slowly enough between dimeric and monomeric states that we could separate them by size exclusion chromatography. We observed differing activity levels of the various oligomerization states of these proteins – both monomers are active, with the MS367 dimer being more active than the monomer, and RE356 displaying opposite results (**Fig. 4**). The *in vitro* data lead us to propose a mechanism where shifts in the monomer:dimer equilibrium could regulate sensor HK enzymatic activity. While demonstrating that monomer:dimer control of RE356 and MS367 activity *in vivo* is outside the scope of this work, we note that comparable mechanisms are widely accepted or suggested for a very broad range of signaling proteins including certain sensor HKs (43), receptor tyrosine kinases (44), and photoreceptors (45-48). In this case, dimerization may be modulated by potential ligand binding in the yet-uncharacterized PAS domains of these HKs, as in other PAS-domain containing signaling proteins (49), or via other mechanisms. Perhaps the relative instability of the dimer state, which forms higher-molecular-weight aggregates over time, could serve as an intrinsic timer specifically on one of the two signaling species to temporally modify a biological response post-activation.

Given the integral role of dimerization for many sensor HKs, we sought to understand the sequence determinants of dimerization with an engineering approach. Using RH376 as a model dimer, we systematically removed combinations of the three insertions it contains compared to the monomeric EL346 (**Fig. 5, S5**), hypothesizing that one or more of these changes could create a monomeric LOV-HK. While each deletion mutant was well-folded as evidenced by binding flavin chromophores and undergoing canonical LOV photochemistry, we found that all three single deletion mutants (RH1, RH2, RH3) remained largely dimeric despite the removal of 5-30 amino acid residues apiece. Remarkably, the combination of two of these deletions to generate the RH1+3 construct substantially monomerized the protein and abolished autophosphorylation activity.

From these studies on RH376, we arrive at the general conclusion that multiple regions determine oligomeric states in this family of LOV-HKs. The use of distributed interfaces for dimerization is consistent with structural studies on the dimeric YF1 engineered sensor kinase (2) showing intermonomer contacts via both the LOV and DHp domains, which are typical dimerization interfaces for LOV photosensors (2, 50) and canonical HisKA histidine kinases (13, 51) respectively. These correspond to our RH1 and RH2 deletions; neither of these changes, nor the RH3 deletion in a large loop near the kinase ATP binding site, were sufficient on its own to generate monomers. Unexpectedly, the RH3 deletion – which removes a large loop near the ATP binding site – affected dimerization when paired with RH1. While this site has not been previously described as being involved in dimerization to the best of our knowledge, we note that ATP itself bridges interactions between multiple domains in two known HWE/HisKA2 HK crystal structures (1, 38), laying some precedent for interactions at or near the nucleotide to be involved in dimerization. Clearly, further structural work is needed to more fully assess how dimerization is encoded among multiple HKs.

The diversity of oligomeric states and function in a family of sensor HKs that we uncovered here opens the door to novel regulatory modes within this family, such as control of dimerization, whether by ligand or inherent thermodynamic stability, or cooperativity. The closely related bacterial chemoreceptor proteins, for instance, form large arrays to increase signaling output through cooperativity (16). Similar use of monomer/dimer transitions and evolution of variable quaternary structure is at the heart of acquiring cooperativity and novel regulatory modes in the classic hemoglobin family of oxygen-binding proteins, which started as ancestral monomers prior to dimerizing by gene duplication events and the acquisition of a relatively small number of mutations (33). Our findings suggest that various oligomeric states within the sensor HK family could allow for different potential for cooperativity, perhaps depending on the input signal or output pathway, or integrating multiple points of control or regulation to be added to the canonical signal transduction pathway of two-component systems.

## Experimental Procedures

### Bioinformatics

Sequences similar to EL346 were identified by a BLAST search with default parameters against the non-redundant NCBI database, using EL346 full-length protein sequence (Uniprot ID: Q2NB77) as a query. EL346 and the best 100 hits were aligned using Clustal Omega (52) and a distance tree was calculated from this alignment using the same software. Graphic display of this tree was accomplished using iTOL (Interactive Tree of Life, http://itol.embl.de/) (53). This tree was used to select HKs for cloning from the strains available at DSMZ (http://www.dsmz.de).

### Cloning, protein expression and purification

Five HKs were selected for biochemical characterization, abbreviated RT349, RH376, ME354, RE356, and MS367 from the combination of their species name (e.g. *Rubellimicrobium thermophilum* = RT) and number of amino acid residues (**Table 1**). DNA encoding these proteins were cloned into a pHis-G*β*1 vector (54), with constructs verified by DNA sequencing. Proteins were overexpressed in *Escherichia coli* BL21(DE3) (Stratagene) in LB as previously described (18). Cells were harvested, resuspended in Buffer A (50 mM Tris pH 8.0, 100 mM NaCl, and 10 mM MgCl_2_), and lysed by sonication. Lysates were centrifuged at 48,000 x g and 4 °C for 45 min. Supernatants were loaded into a Ni^2+^ Sepharose affinity column (GE Healthcare) and the His6-G*β*1 tagged protein was washed with 4 column volumes of Buffer A supplemented with 15 mM imidazole and eluted with 4 column volumes of Buffer A supplemented with 250 mM imidazole. Eluted proteins were exchanged into Buffer A by dialysis, then fusion tags were cleaved by incubation with His_6_-TEV protease overnight at 4 °C. The tag-less proteins were separated from the tags and His_6_-TEV protease by Ni^2+^ affinity chromatography and were further purified by size-exclusion chromatography on a HiLoad 16/600 Superdex 200 column (GE Healthcare) equilibrated with HSEC buffer containing 50 mM HEPES pH 7.5, 100 mM NaCl, 10 mM MgCl_2_, and 0.5 mM DTT. For light-sensitive proteins, all purification steps were performed under dim red light. Concentrations were determined from the theoretical absorption coefficient, ε_280_ for PAS-HKs, calculated from the sequence using the ExPASy ProtParam server (55), and ε_446_ = 11,800 M^-1^ cm^-1^ for flavin-containing proteins.

RH376 deletion mutants were generated as follows: RH2 and RH3 were ordered as synthetic constructs from Genewiz (South Plainfield, NJ) and inserted into the same pHis-G*β*1 vector as the WT protein. RH1 and RH1+3 mutants were generated by PCR using an N-terminal primer lacking the sequence to be deleted using WT and RH3 plasmids as templates, respectively.

### Absorbance and dark state reversion kinetics

UV-visible absorbance spectroscopy measurements were acquired using a Varian Cary 50 spectrophotometer at 24 °C with a 1 cm path length quartz cuvette. First, a UV-visible spectrum in the range 250-550 nm was acquired in light and dark conditions for each LOV-HK. For dark measurements, all steps were performed under dim red light. For light measurements, the samples were illuminated for 1 minute using a blue LED panel (225 LED bulbs in 30.5 × 30.5-cm panel, 13.8 W, 465 nm maximum illumination wavelength; LEDwholesalers [Hayward, CA]) prior to each scan. To measure dark state recovery, each LOV-HK sample was illuminated under a blue LED panel for 1 minute and then transferred to the spectrophotometer. A_446_ of the samples in HSEC buffer with 1 mM ATP was monitored at 5-min intervals for 400-800 min at 24 °C. Time constants *τ* were determined by fitting measurements to a monoexponential decay using the following equation:

Abs = A_inf_ - (A_inf_ -A_o_)e^-t/^*^τ^*, where A_inf_ = A_446_ at infinity, A_o_ = A_446_ initial, and t = time Raw data were fit and plotted using RStudio with the nls function; plots were edited in Adobe Illustrator CS6.

### SEC-MALS

All samples and buffers were filtered with a 0.1-µm pore size filter before use. ATP was added to each HK sample (∼15 μM, 400 μL) to a final concentration of 1 mM. Samples were then injected at a 0.4 mL/min flow rate onto a Superdex 200 GL 10/300 (or in the case of Figure S4, Superdex 200 Increase GL 10/300) SEC column (GE Healthcare) using HSEC buffer with 1 mM ATP. For lit experiments, samples were illuminated with a blue LED panel for 1 min prior to injection and the blue LED panel was faced toward the column during the run. For dark experiments, all steps were performed under dim red light. Samples were detected post-elution by the inline miniDAWN TREOS light scattering and Optilab rEX refractive index detectors (Wyatt Technology [Santa Barbara, CA]). FPLC runs were performed at 4 °C, while MALS and refractive index measurements were at 25 °C. Data analysis and molecular weight calculations were performed using the ASTRA V software (Wyatt Technology). Raw data were plotted using RStudio. Assignments of oligomeric state are based on a qualitative, five-bin scale, with proteins considered to be “dimeric” or “monomeric” when their MALS-calculated mass falls within ±5% of the sequence-derived molecular weight for either state. Between these ranges are three bins of equal size: “mostly dimer”, “monomer/dimer mixture”, and “mostly monomer.”

### Autophosphorylation assays

Autophosphorylation was measured as described previously (1, 18). A minimum of three trials were conducted for each protein under each condition. Proteins were added to a reaction buffer containing 50 mM Tris, 100 mM NaCl, 5 mM MnCl_2_, and 1 mM DTT with pH 8.2. Protein concentrations were confirmed by analysis of dilution series on Coomassie-stained SDS-PAGE. Final reaction concentrations ranged from approximately 10 to 35 micromolar as listed in the table below:

| *Protein* | *Concentration (μM)* | *Protein* | *Concentration (μM)* |
| --- | --- | --- | --- |
| EL346 | 16.1 | RE356 monomer | 9.6 |
| ME354 | 11.6 | RE356 dimer | 8.5 |
| RT349 | 28.6 | RH1 | 12.4 |
| RH376 WT | 14.3 | RH2 | 30.4 |
| MS367 monomer | 10.7 | RH3 | 34.7 |
| MS367 dimer | 11.1 | RH1+3 | 9.1 |

A mixture of unlabeled ATP and 10 *μ*Ci [*γ*-^32^P] ATP was added to each mixture to initiate the reaction (final ATP concentration of 200 μM). Aliquots were removed at time points of 0.5, 1, 2, 4, 8, 6, and 32 min and placed into a 4x SDS-gel loading buffer to quench the reaction. For dark measurements, all steps were performed under dim red light. For light measurements, the samples were illuminated with a blue LED panel just prior to and throughout the course of the experiment. Samples were subjected to SDS-PAGE analysis; dried gels were exposed for 1hr - overnight and bands were visualized by phosphorimaging. Band intensities were measured using Fiji (56, 57); initial velocity values were calculated in Microsoft Excel from the linear region of the curve and were plotted in RStudio.

## Data availability

All data for this work are contained in this manuscript.

## Supporting information

This article contains supporting information, including Figures S1-6, Table S1, and accompanying literature citations.

## Acknowledgements

The authors would like to acknowledge Bright Shi for generating the RH1+3 construct, as well as Matthew Cleere, Zaynab Jaber, Jaynee Hart, and other Gardner lab members for fruitful discussions.

## Funding and additional information

This work was supported by NIH grants R01 GM106239 (to K.H.G.) and F32 GM1119311 (to I.D.). The content is solely the responsibility of the authors and does not necessarily represent the official views of the National Institutes of Health.

## Abbreviations

HK: histidine kinase
LOV: Light-Oxygen-Voltage
PAS: Per-ARNT-Sim
TCS: two-component system
RR: response regulator
DHp: dimerization histidine phosphotransfer
CA: catalytic ATP-binding
SEC-MALS: size exclusion chromatography multi-angle light scattering.

