## Supporting Information for "Diversity of function and higher-order structure within HWE sensor histidine kinases"

**Conservation of function with diversification of higher-order structure within sensor histidine kinases**

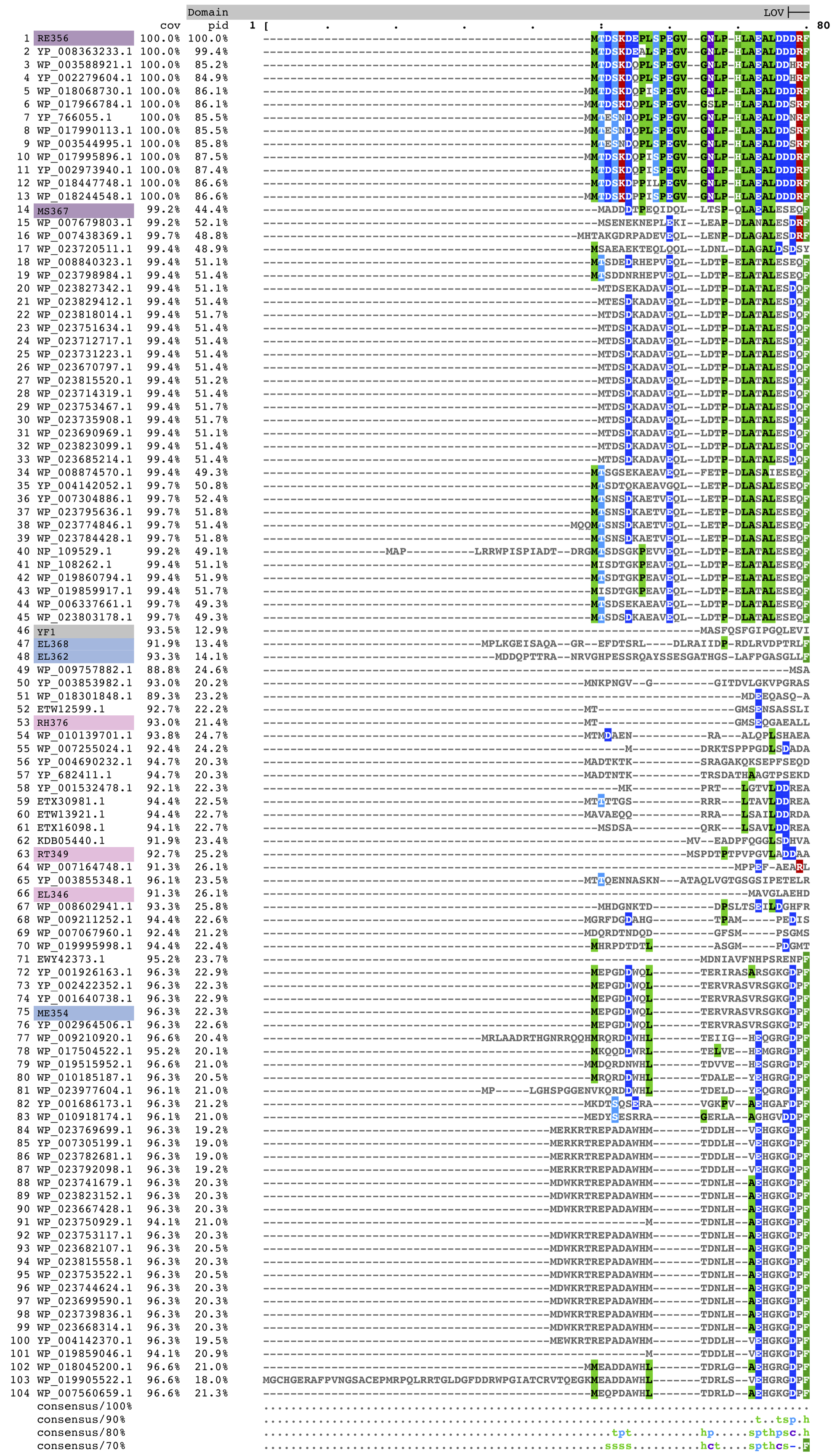

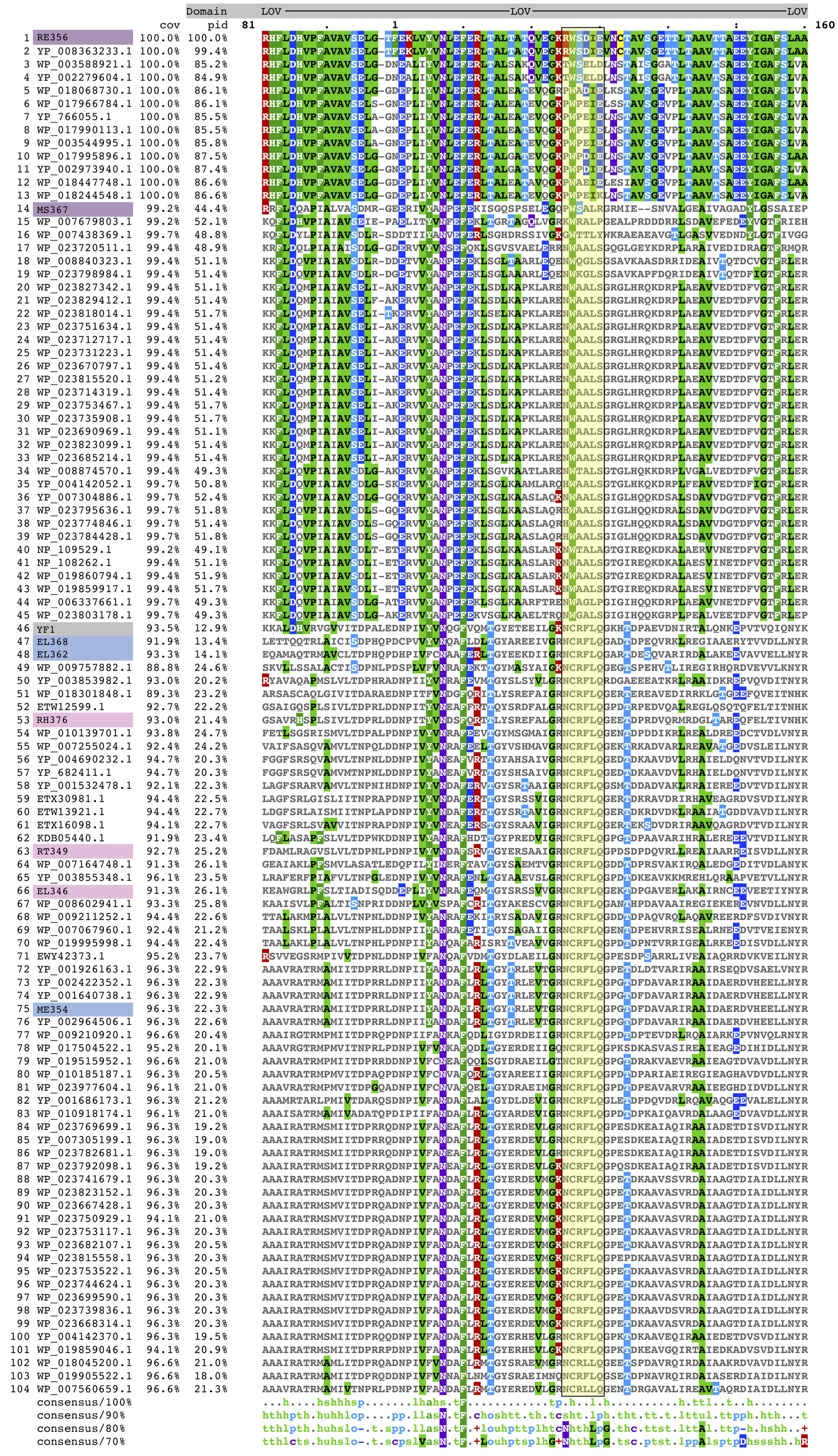

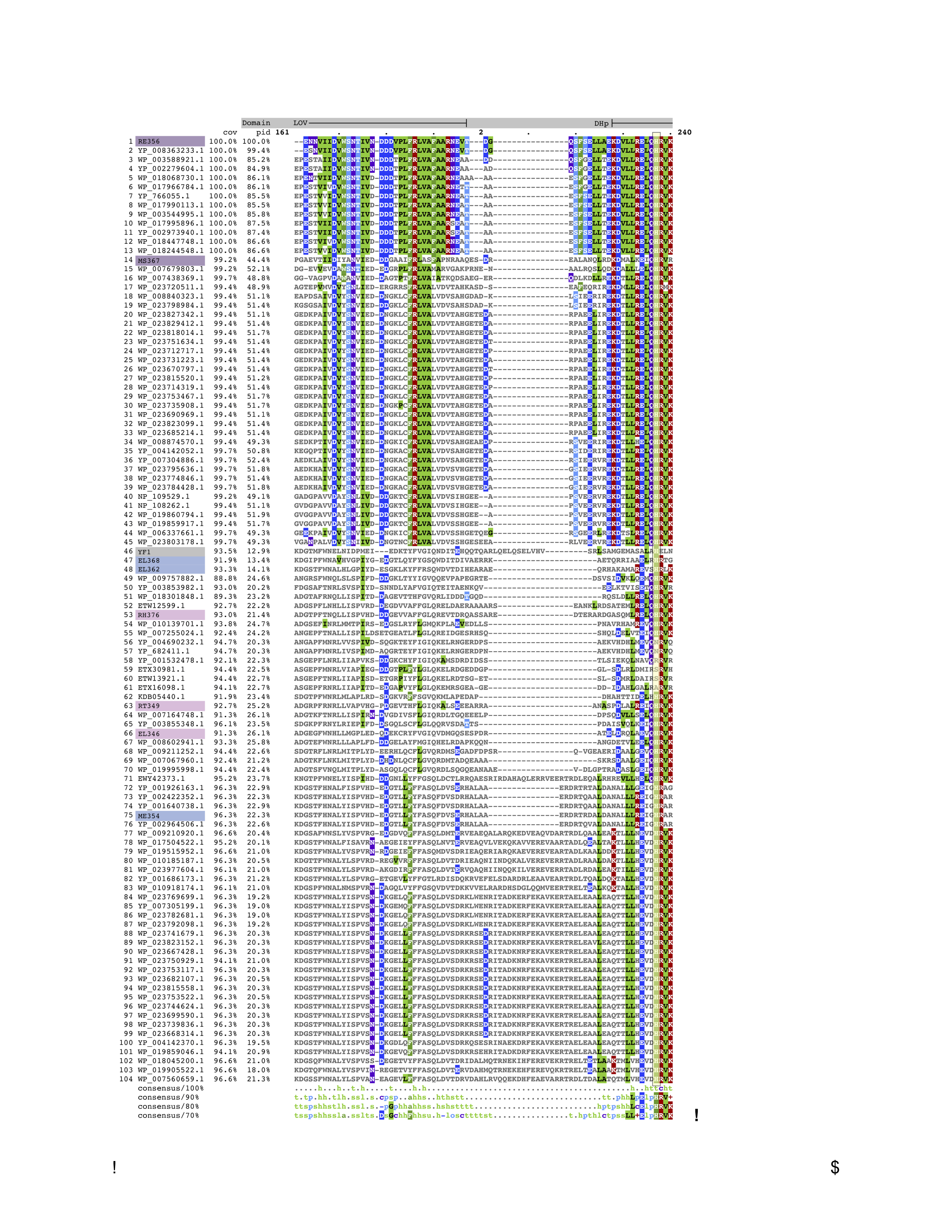

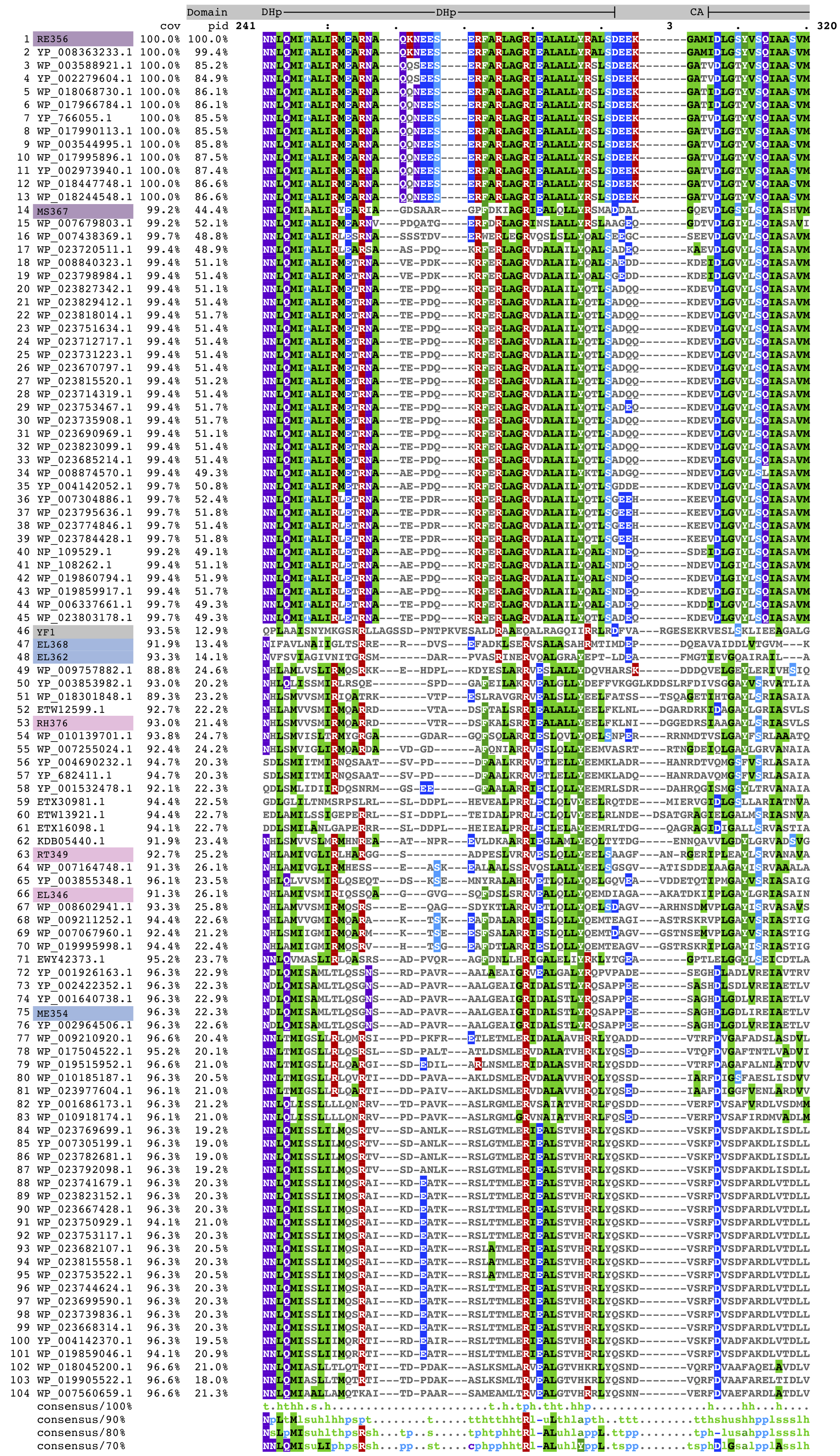

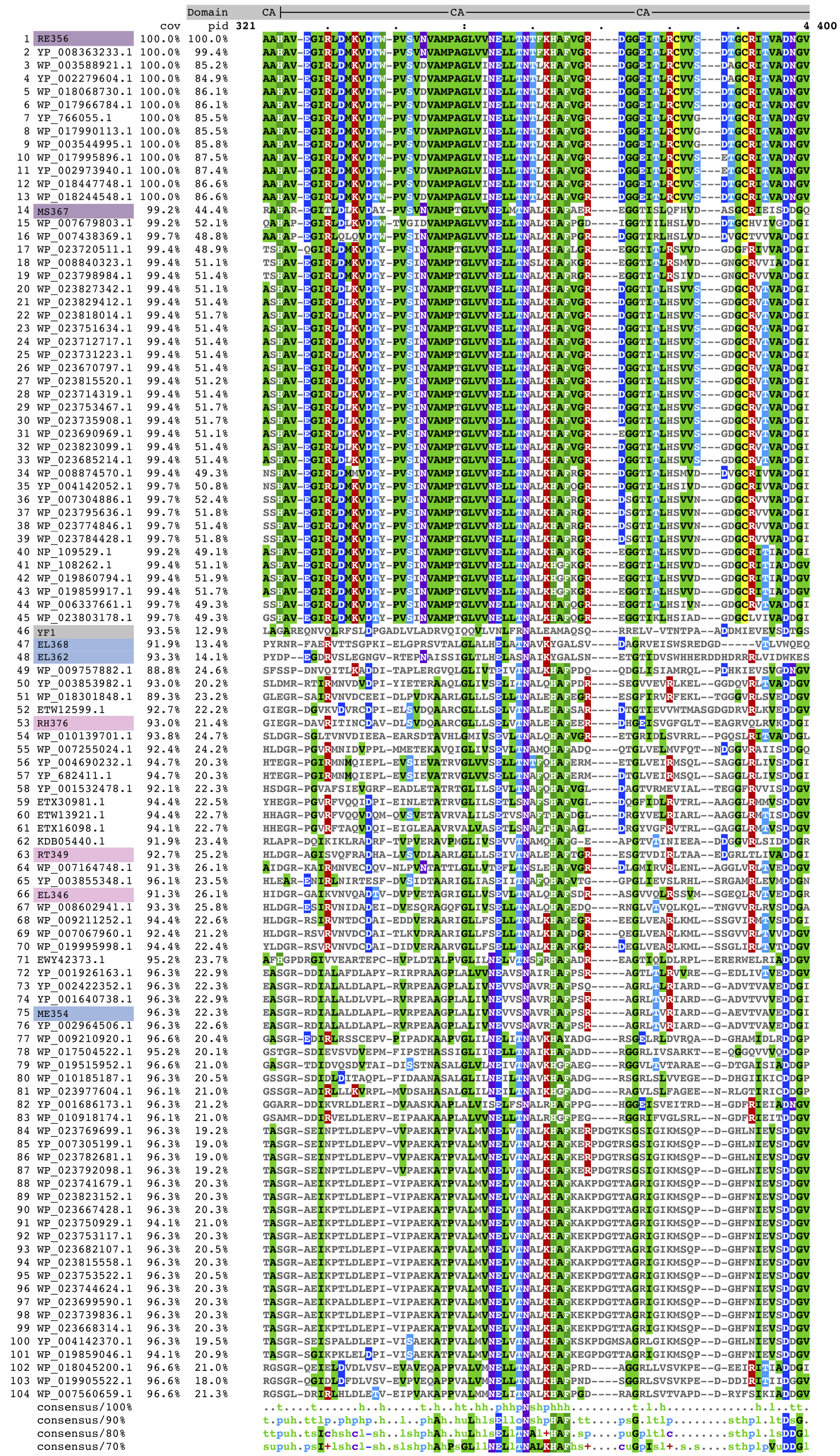

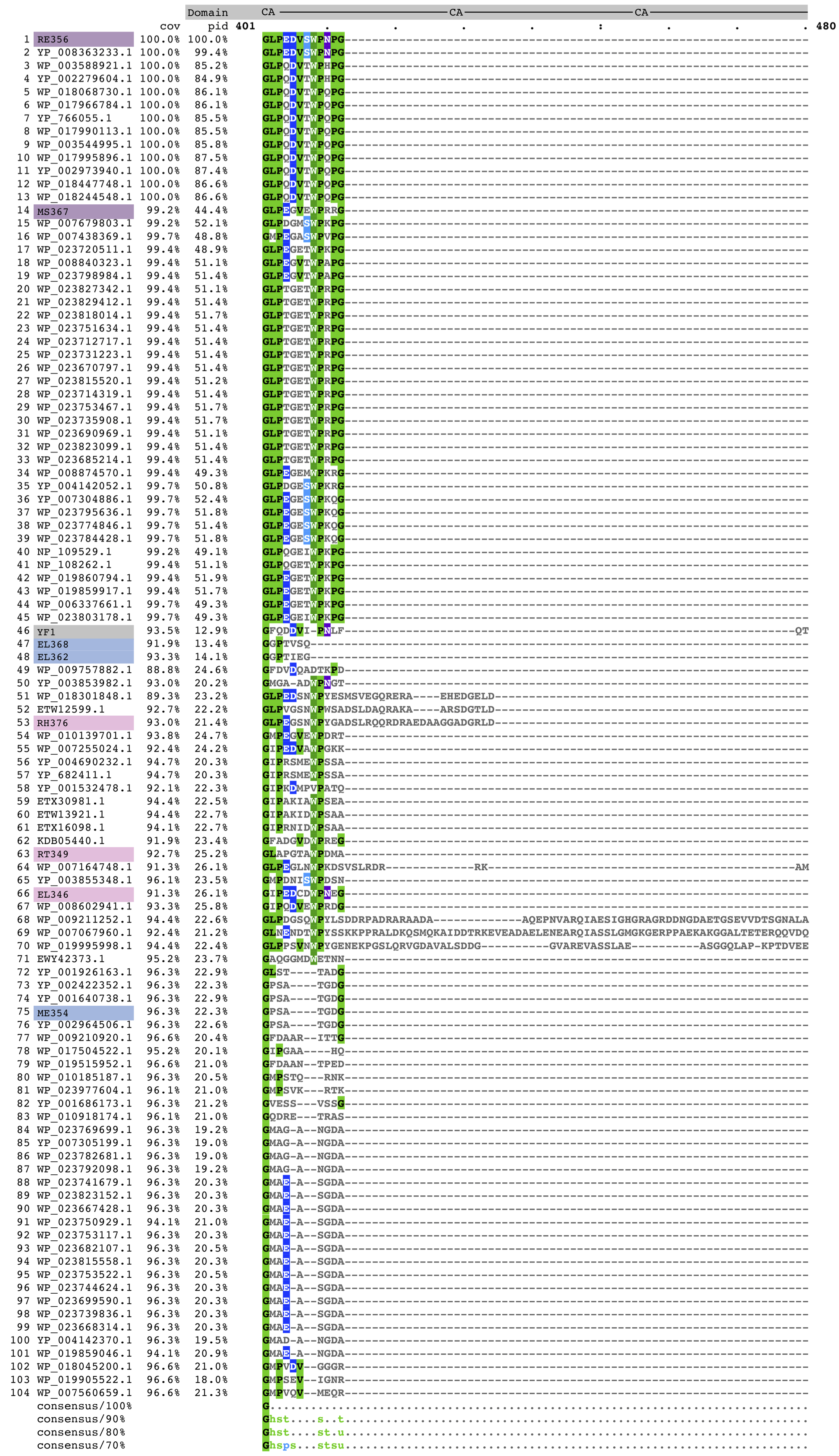

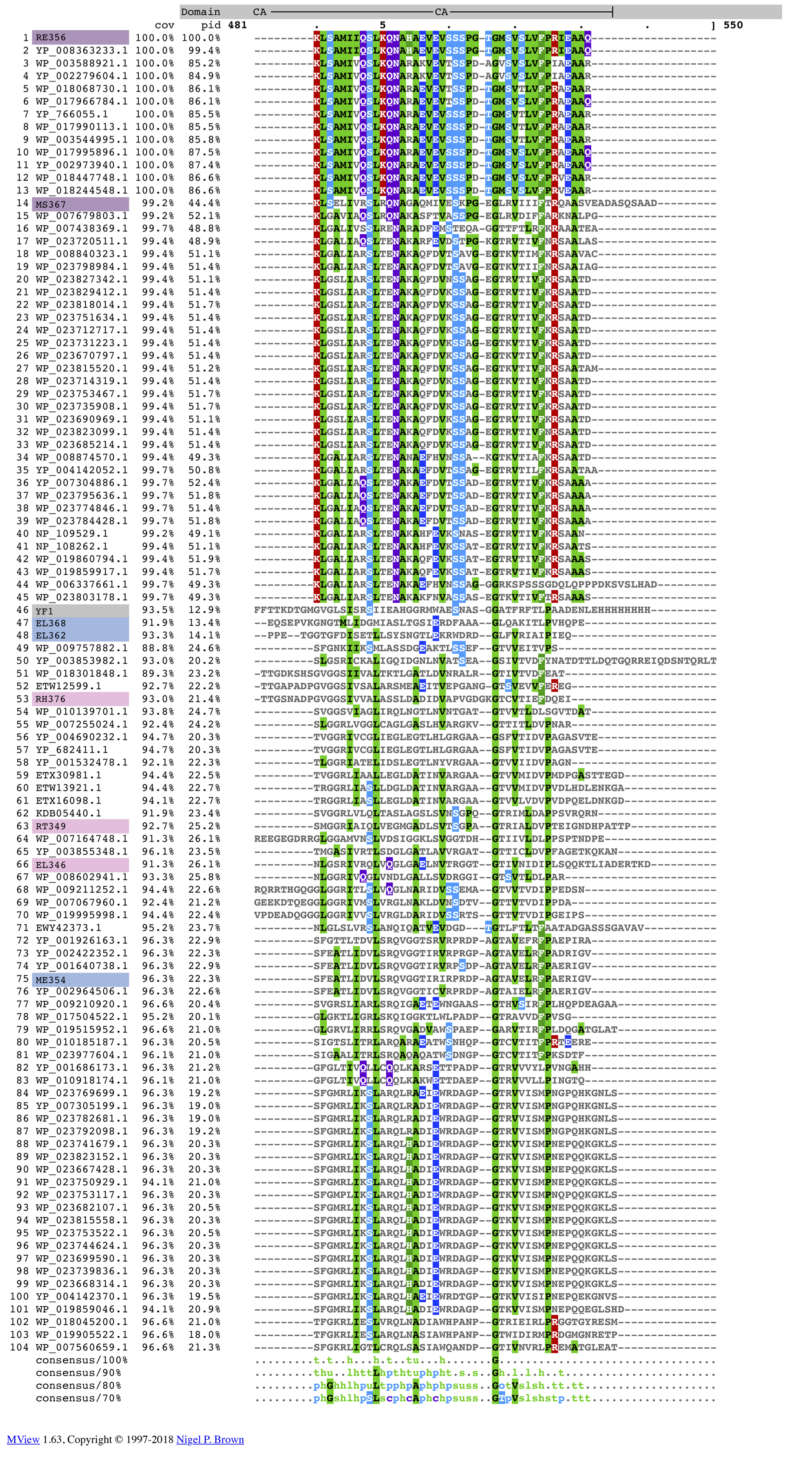

**Figure S1: Multiple sequence alignment of top 100 BLASTp EL346 homologues.** Multiple sequence alignment was generated with Clustal Omega (1) and visualized in MView (2). All sequences are labeled by GenBank accession numbers, while the HKs focused on in this study are labeled with the names they are referred to by in this paper and highlighted with the color corresponding to their clade (**Table 1**). Domain architecture of EL346 is shown on top in grey. Percent identity to EL346 sequence is shown in the “pid” column. The phosphoacceptor histidine region (pg. 3) and characteristic “NCRFLQ” LOV sequence (pg. 2) (3) are highlighted in yellow.

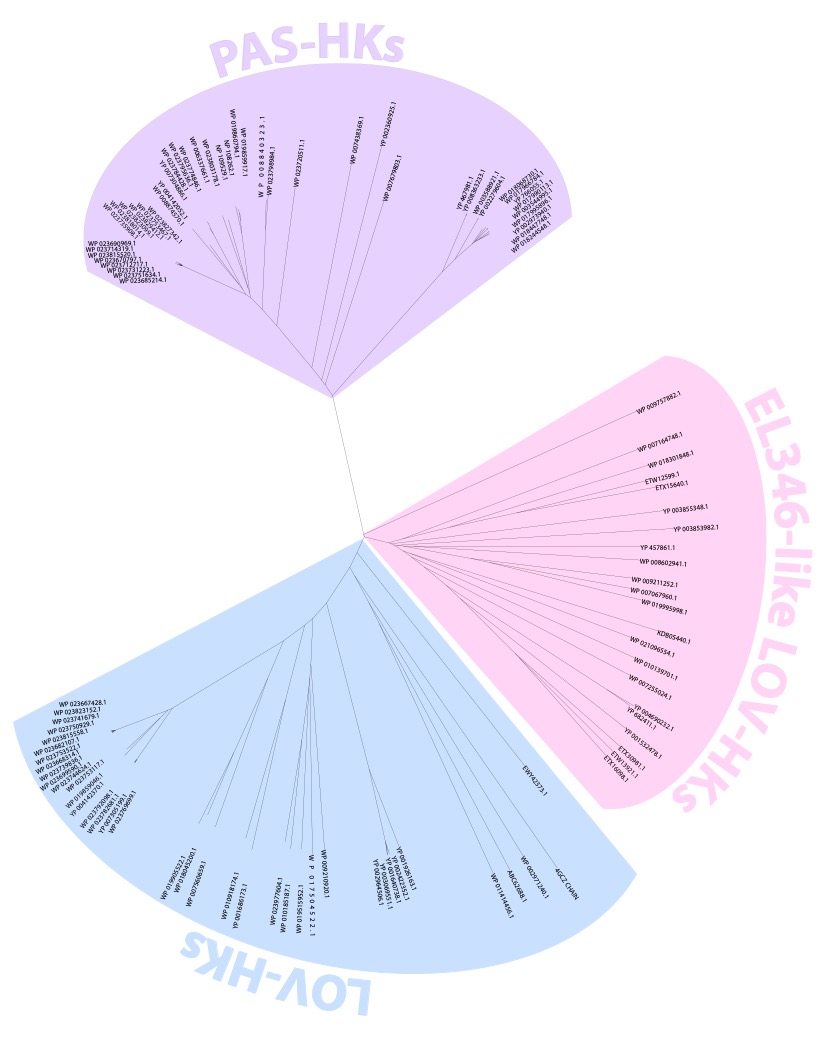
**Figure S2: MSA Tree of EL346 Homologs.** NCBI reference numbers for each of the 100 EL346 homologs shown in Figure 1 are listed here.

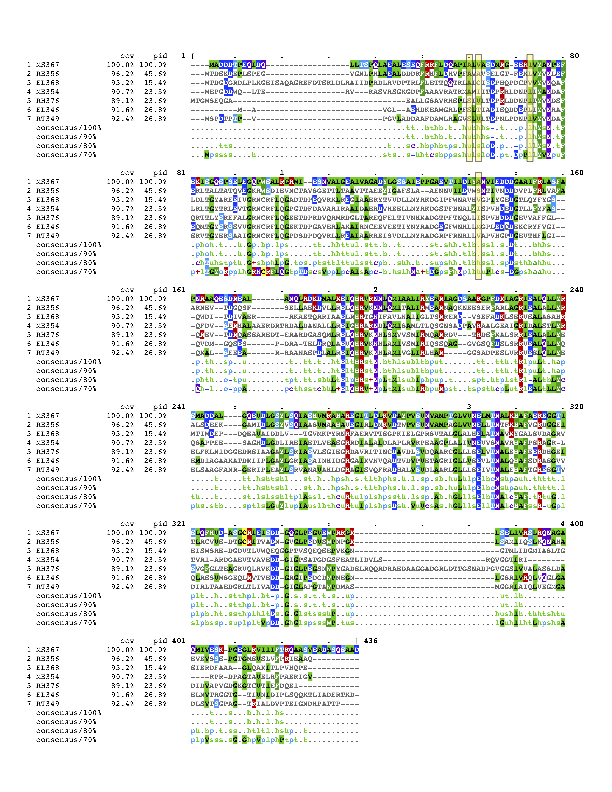

**Figure S3: ME354 and RT349 primary sequences include “slowing mutations” that impact the rate of LOV domain dark state reversion.** Multiple sequence alignment was generated with Clustal Omega (1) and visualized in MView (2). Mutations (4) at positions #19, 21, 32, and 101 in EL346 are highlighted in yellow.

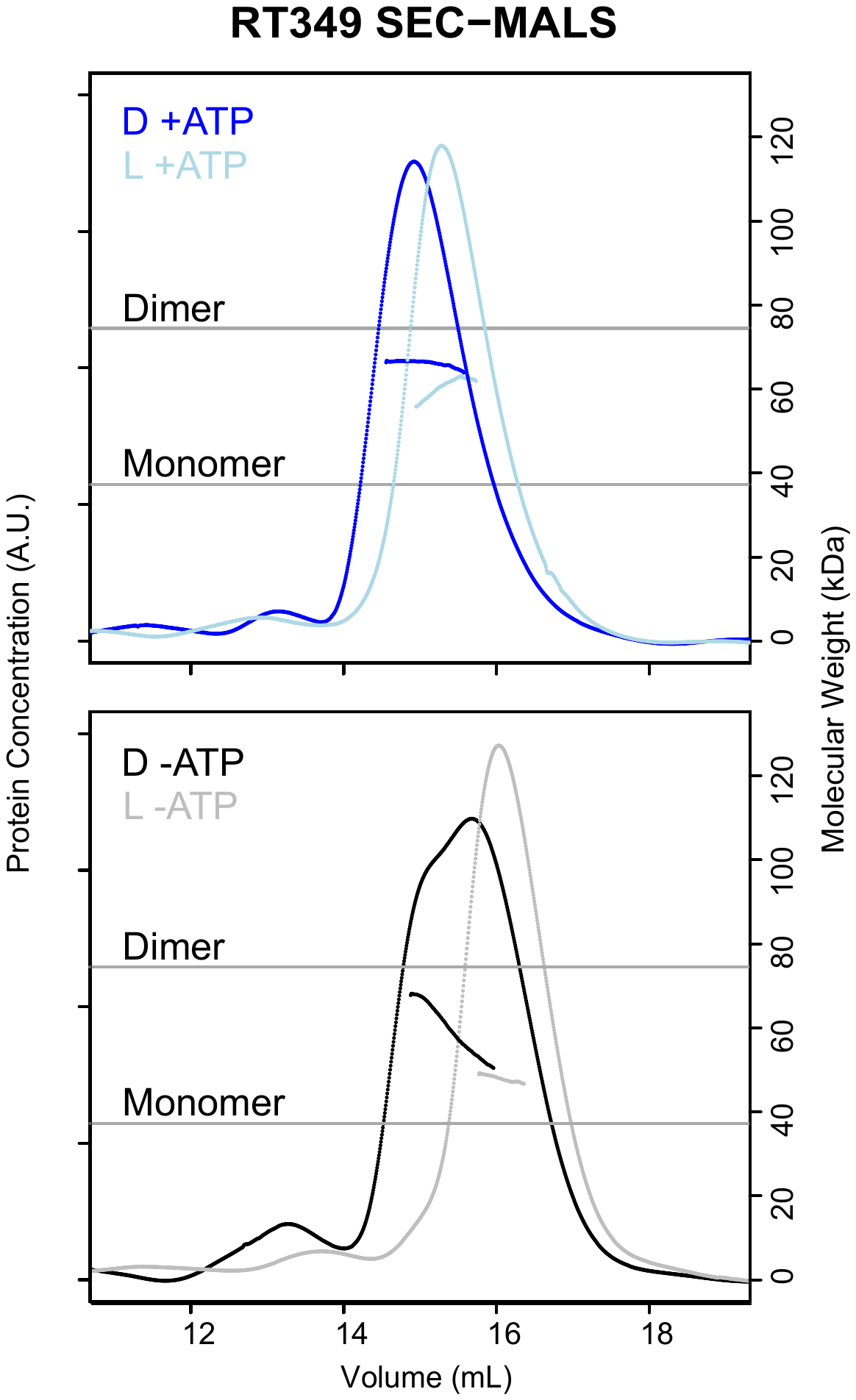

**Figure S4: RT349 elution peak shift is dependent upon the presence of ATP.**

**Top)** Shift of SEC-MALS peak in the presence of 1 mM ATP is reproducible on the higher-resolution Superdex 200 Increase 10/300 GL SEC (GE Healthcare). **Bottom)** SEC-MALS of RT349 in the absence of ATP reveals two additional shifted peaks indicating a continuum of oligomeric and conformational states and a role for ATP in the conformational changes of RT349.

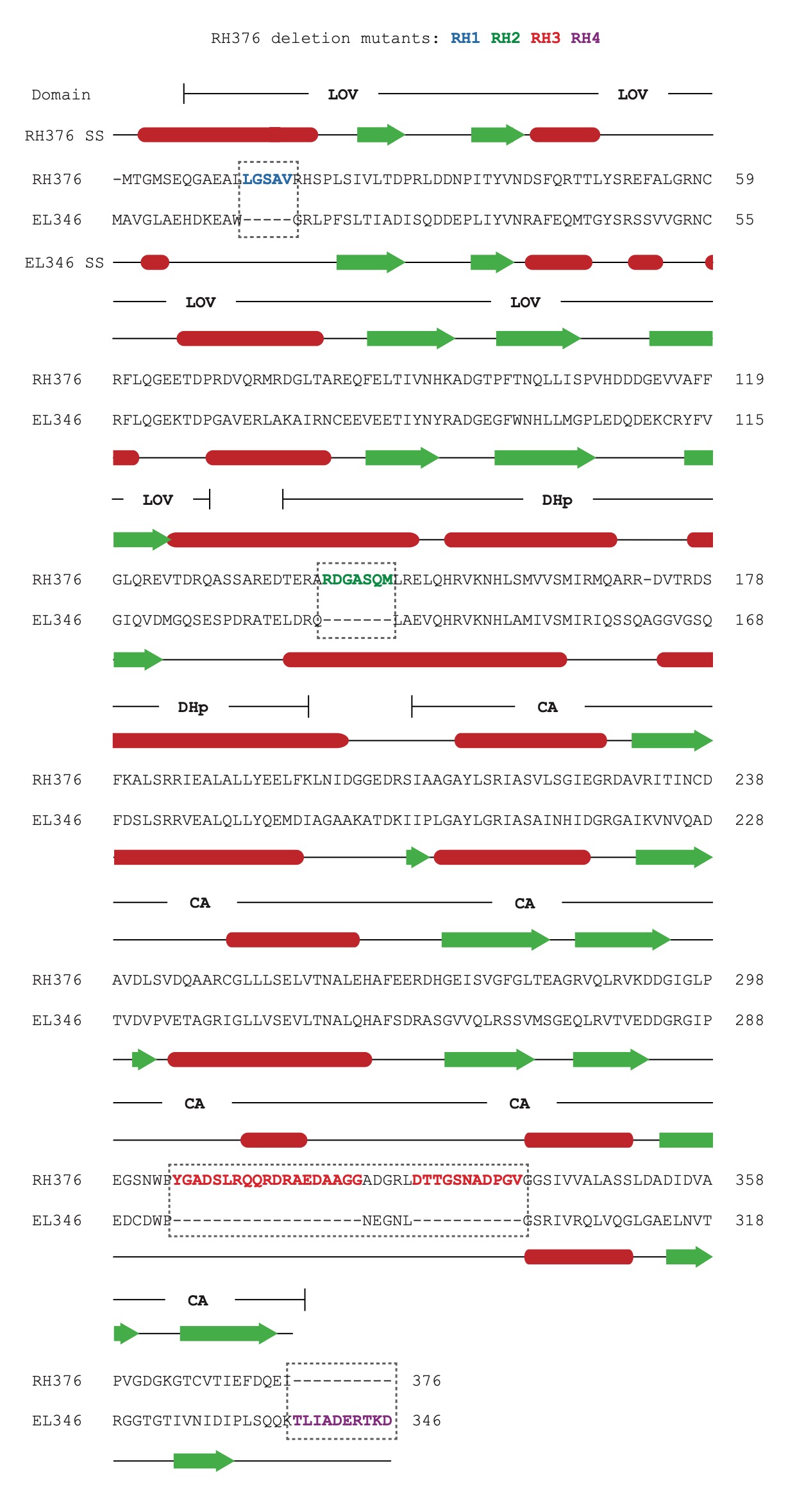

**Figure S5: Pairwise alignment of EL346 and RH376 reveals “gaps” in EL346 sequence.** The pairwise alignment of EL346 and RH376 sequences was performed by MUSCLE (5). Regions of difference between EL346 and RH376 termed RH1-4 are highlighted with dashed boxes. Secondary structure prediction by JPred (6) (red = helices, green = sheets) for RH376 and from the 4R3A PDB entry for EL346 (7) are shown above and below the sequences, respectively. Domain locations (LOV, DHp, and CA) are shown above the amino acid sequences.

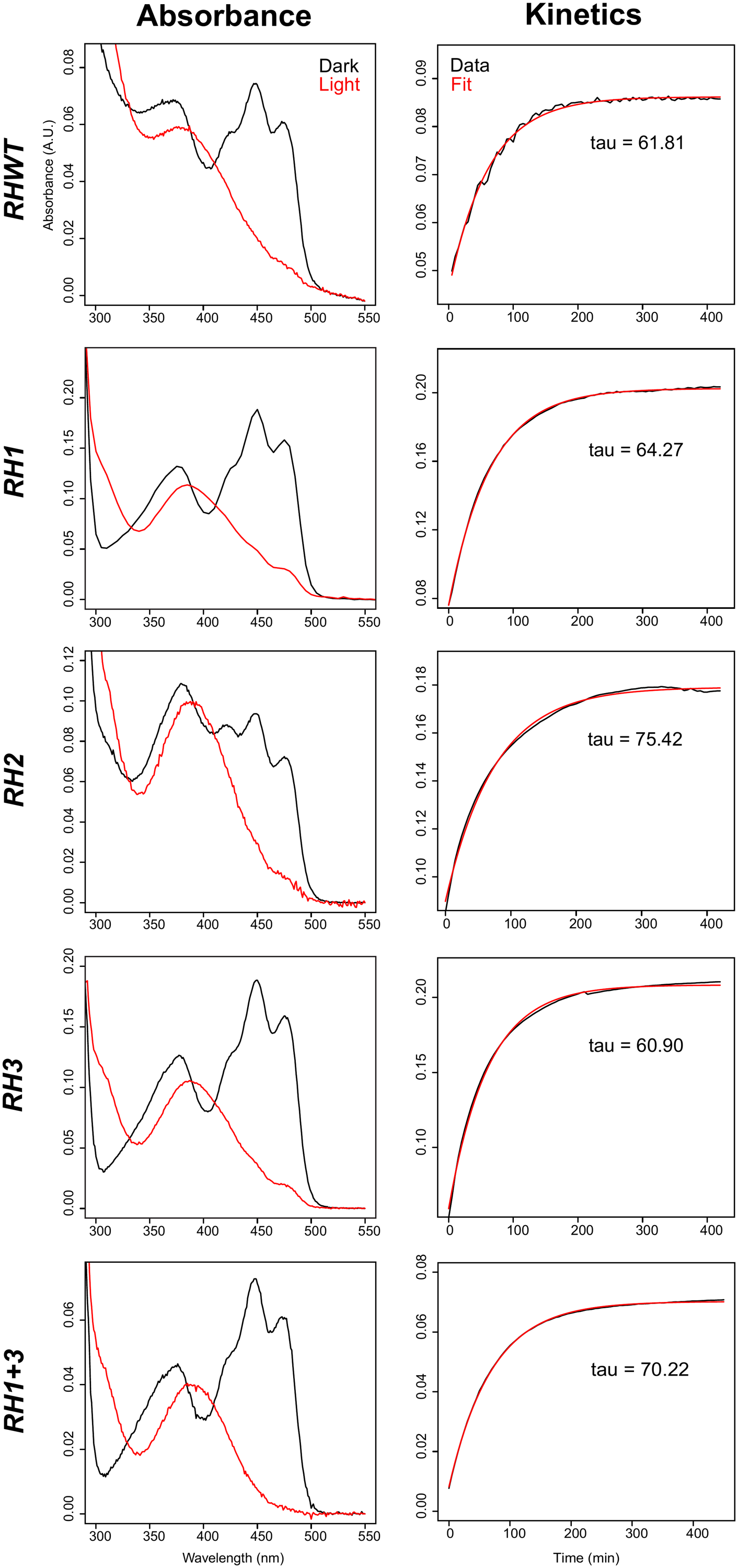

**Figure S6: RH376 mutant UV/visible absorbance and dark state reversion kinetics show that mutants are light sensitive with recovery times comparable to the WT. a)** Absorbance of dark and light states along with **b)** dark state reversion kinetics show that RH376 WT and mutants are light sensitive. Based on tau values extracted from a single exponential decay fit, the photocycle of these mutants is comparable to the WT.

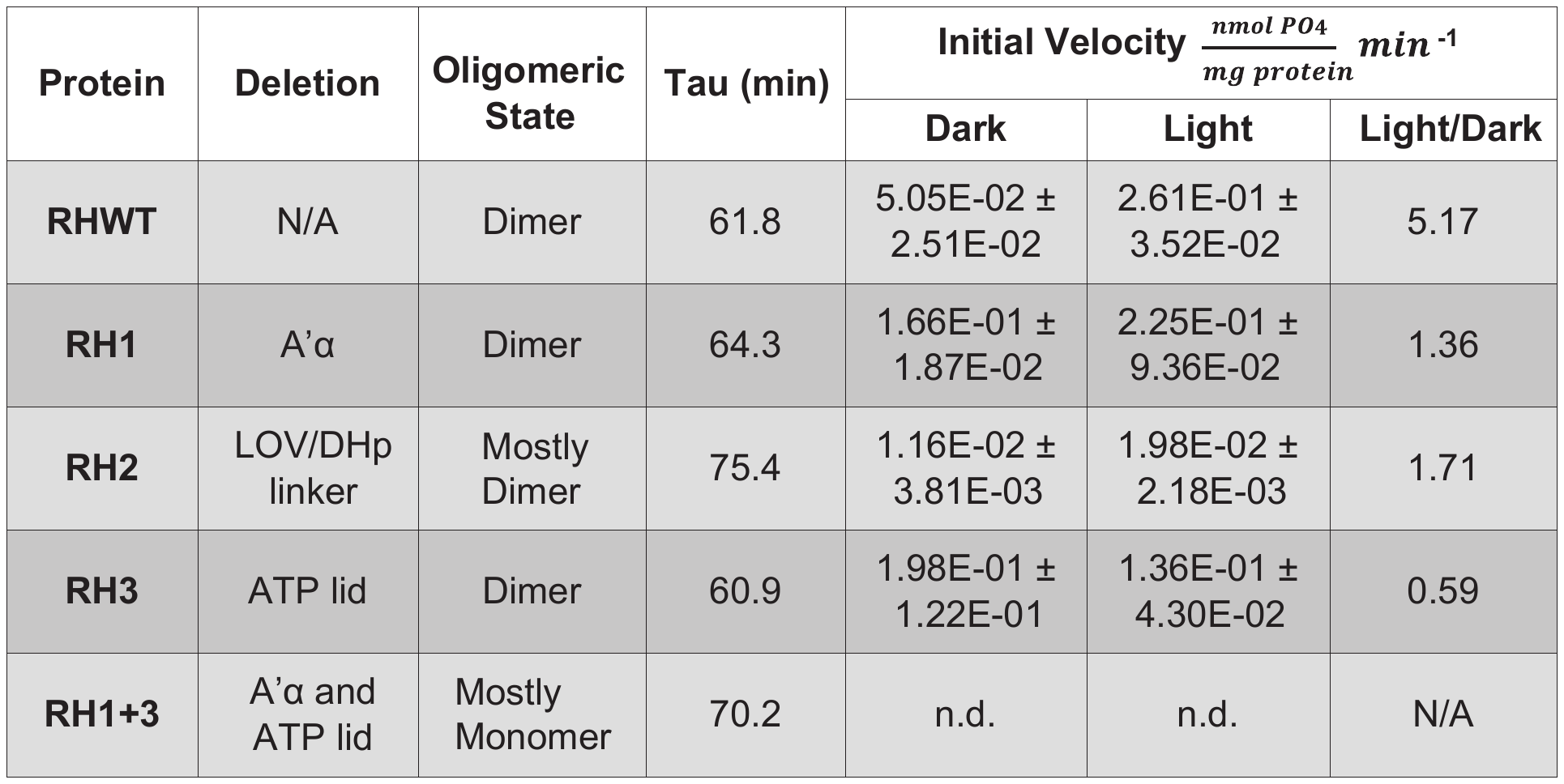

**Table S1: RH376 Deletion Mutant Measurements.** Mass data were collected using SEC-MALS (n=1), and oligomeric states are defined using a qualitative five-bin scale. Tau represents the time constant of the thermal reversion of the light state as fit to a first-order exponential decay (Fig. S6). Initial velocities were calculated from the linear region of autophosphorylation activity curves. Error is represented by the percent bias between trials (n=3) of the same conditions. The ratio of light to dark initial velocities represents the degree of light activation that each protein undergoes.
